# Interaction of involuntary and executive attention during development

**DOI:** 10.64898/2026.07.19.739397

**Authors:** Ursula Schöllkopf, Andreas Widmann, Aurélie Bidet-Caulet, Nicole Wetzel

## Abstract

Attentional control requires the fine-tuned interaction of attentional networks supporting alertness, orienting and higher-level executive control. This study examined the interaction of executive attention, comprising inhibition, and involuntary orienting towards unexpected deviant sounds in children (6–8-years, N=30), adolescents (10–12-years, N=39) and adults (18–34-years, N=35). An auditory equiprobable Go/Nogo-Oddball paradigm was employed to investigate executive (Go/Nogo) and involuntary (Oddball) attention and their interaction. Event-related potentials (ERP) in the EEG, pupil dilation and behavioral data were analyzed using Bayesian statistics.

Deviant sounds evoked an involuntary attentional orienting reflected by the ERP component P3a and decreased performance in all age groups. The distraction effect diminished with age. Pupil dilation in response to deviant and target sounds were modulated by involuntary and executive attention but showed no interaction. In the EEG, frontal Nogo-N2 and Nogo-P3 effects, which reflect response inhibition, appeared in standard trials but not in deviant trials. This interaction suggests that response inhibition is reduced by involuntary attention orienting. While all age groups showed similar amplitudes of the Nogo-N2 effect, the subsequent Nogo- P3 effect, which has been associated to motor inhibition, was absent in the 6–8-year-olds. Age differences observed in latencies of the Nogo-N2 effect but not in the latencies of the Nogo- P3 effect (between adolescents and adults) indicate distinct developmental trajectories of the response inhibition mechanisms underlying the Nogo-N2 and Nogo-P3 effects throughout childhood and adolescence.

The present data provide novel insights into the interaction between executive and orienting attention networks in the auditory modality during development.

## 1 Introduction

Attentional control ensures the efficient processing of goal-relevant information by flexibly allocating attentional resources to task-relevant information while minimizing the processing of irrelevant information (Markant & Amso, 2022). According to the influential Posner model (1990), attention includes three primary networks: (1) The *alerting network*, which is necessary to achieve and maintain an alert state. (2) The *orienting network* is responsible for the voluntary or involuntary orienting of attention towards a task-relevant event or a salient stimulus, respectively. (3) The *executive network*, which manages higher-level, voluntary control processes (executive attention) including target detection, selection and inhibition (for review see Posner & Rothbart, 2023).

In everyday life, these networks interact and complement each other. In educational settings, successful attentional control enables students to voluntarily stay focused, resist involuntary distractions and process information effectively leading to better academic performance and classroom behavior. The balance between voluntary and involuntary attention, and their interaction, are essential for successful attention control but has been less understood from a developmental perspective. In the literature, processes attributed to different attentional networks are often studied separately. Few studies investigating the interaction of attentional networks focused on the visual domain and used behavioral measures (Callejas et al., 2004; Fan et al., 2002; Pozuelos et al., 2014). In the auditory domain, the interplay between those networks and particularly the underlying brain dynamics are not yet sufficiently understood, especially during development. The present study aims to close this gap and examines the interaction between the orienting and executive attention networks by combining distraction and inhibition in a newly developed auditory Go/Nogo-task, in children (6–8y) and adolescents (10–12y) and in an adult group, using pupillometry and EEG.

### 1.1 Involuntary orienting of attention

Involuntary orienting of attention describes the capture of attention by task-irrelevant, but potentially important events outside the current focus. This can lead to impaired performance in the current task (distraction effects). Versions of the auditory oddball paradigm were used to study involuntary orienting of attention, where participants are presented with a stream of frequent *standard* sounds and infrequent *deviant* or *oddball* sounds. The latter involuntarily capture attention leading to impaired behavioral performance in adults (e.g. Schröger and Wolff, 1998) and children (for review see Wetzel & Schröger, 2014). Those distraction effects are typically larger in children and decrease through early and middle childhood (Volkmer et al., 2022; Wetzel et al., 2019) along with the maturation of underlying brain networks. This maturation begins in primary sensory regions and gradually progresses through secondary and multimodal regions with the prefrontal cortex being one of the last areas to fully mature during young adulthood (e.g. Gogtay et al., 2004; Toga et al., 2006). In the EEG, the orienting response, encompassing the involuntary orienting of attention and an increase in arousal, has been associated to a component in the event-related potential (ERP) with a fronto-central topography peaking around 250-300 ms after sound onset, the P3a (Polich, 2007; Masson, 2019). Moreover, a larger pupil dilation response (PDR) has been observed to deviant sounds compared to standard sounds (Bonmassar et al., 2020; Eckstein et al., 2017; Widmann et al., 2018). Pupil size is an indirect measure of the locus coeruleus norepinephrine system (LC-NE) activity (Einhäuser, 2017; Murphy et al., 2014). As the main source of NE in the cortex, LC has a considerable influence on signal processing in respective target areas and is thus crucial for cognitive functions including attention (Aston-Jones & Cohen, 2005; Joshi et al., 2016; Poe et al., 2020).

### 1.2 Executive attention

Executive attention, which includes target processing and inhibition, can be investigated using the Go/Nogo task where subjects must respond to one stimulus (Go) and inhibit the response to the other (Nogo). Typically, rare Nogo stimuli are presented in a stream of frequent Go stimuli, which creates a prepotent response tendency to act. On the behavioral level, a high rate of false alarms (FA) reflects poor response inhibition while hit rate and reaction time (RT) give insight into target processing (for meta-analysis see Wright et al., 2014). It has been shown that reaction times and error rates are decreasing with age from early childhood to adulthood (for review see Weiss & Luciana, 2022). In the EEG, the ERP exhibits larger amplitudes in Nogo than Go trials in the time range of N2 (more negative) and P3 (more positive) components which are suggested to reflect inhibitory processes (for review see Cheng et al., 2019; Pires et al., 2014). Following Smith et al. (2008) we use the term *Nogo-N2 effect* and *Nogo-P3 effect* to refer to these increased amplitudes observed in Nogo relative to Go trials at frontal electrode locations in the N2 (200−300ms in adults) and P3 time-windows (300−500ms in adults). The Nogo-N2 effect has been associated with response inhibition or response conflict-monitoring. The Nogo-P3 effect has been discussed to reflect cognitive and motor inhibition (Bruin & Wijers, 2002; Smith et al., 2008). In addition, Go stimuli (targets) evoke the P3b component at parietal leads that has been associated to various processes including the allocation of attention resources (Kahneman, 1973; Polich, 2007).

However, there has been considerable debate in the literature regarding the specific cognitive processes linked to the Nogo-N2 effect and Nogo-P3 effect (e.g. Ouyang et al., 2013; Pires, 2014), which can at least in part be attributed to their proximity in latency leading to partially overlapping components. Importantly, response conflict and the requirement to inhibit a response to a Nogo stimulus are also highly dependent on the task (e.g., Go/Nogo task, continuous performance test, Stop signal task), and the target probability (Bruin & Wijers, 2002; Nieuwenhuis & Yeung, 2003; Hoyniak, 2017).

Developmental studies mostly presented Go and Nogo trials with different probabilities. A meta-analysis of such studies specified that N2 amplitudes in Nogo trials, but not in Go trials, decreased with age (Hoyniak, 2017). This supports the hypothesis that the Nogo-N2 effect reflects maturation in brain areas involved in response inhibition (Hoyniak, 2017). The few studies on the Nogo-P3 effect reported increasing amplitudes with age and occasionally a topography shift from parietal towards frontal regions (Hämmerer et al., 2010). These findings suggests an ongoing development of inhibitory control as frontal areas are maturing until young adulthood (Hoyniak, 2017; Johnstone et al., 2005, 2007). However, the use of different paradigms to measure inhibitory processes leads to inconsistencies in the results of developmental studies on the Nogo-N2 and Nogo-P3 effect. For example, during a continuous performance test with infrequent cued Go trials (CPT-AX), a frontal Nogo-P3 effect has been observed in adults but not in 6−7-children and not or just emerging in 9−10-year-old children, indicating increasing inhibition abilities throughout middle childhood (Johnstone et al., 2005; Jonkman, 2006; Jonkman et al., 2003). In contrast, a frontal P3 in Nogo trials has been observed in 8−13- and 9−11-year-olds performing an unwarned equiprobable Go/Nogo task (Barry et al., 2014, 2018).

### 1.3 This study and hypotheses

We aimed to investigate the interplay between involuntary attention and executive attention during development. We combined an auditory oddball distraction paradigm originally introduced by Schröger & Wolff (1998) with an equiprobable Go/Nogo task. Deviant oddball sounds evoked an involuntary orienting of attention while the Go/Nogo task primarily required executive attention abilities. We included two pivotal age groups: 6−8 years, characterized by strong distractibility, and 10−12 years, marked by reduced inhibition abilities (e.g. Hoyer et al., 2021; Johnstone et al., 2005; Wetzel et al., 2019). We measured responses both at the behavioral (reaction times, RT, accuracy) and the physiological levels via pupillometry and EEG. Importantly, to minimize overlap in EEG responses, the Go/Nogo information was provided 200ms after sound onset.

Distraction was measured by longer RTs for deviants compared to standard sounds and was expected to be decreasing with age (Weiss & Luciana, 2022; Wetzel et al., 2019). We expected larger pupil dilation responses to deviants and targets compared to standards in all age groups (Zekveld et al., 2018). Since pupil data provided evidence against an interaction between involuntary and executive attention, details on age effects are provided in the supplementary information (S2.2, Fig S2). In the EEG, we expected inhibition mechanisms, reflected by the Nogo-N2 and Nogo-P3 effects, to be impaired by deviant sounds. Due to increased distractibility and immature inhibition abilities in children and teens compared to adults, we hypothesized that involuntary attention would have a greater influence on inhibition processes in younger than older age groups.

## 2 Methods

### 2.1 Participants

119 participants took part in this experiment. The data of two older and 13 younger children were excluded due to > 60 % errors (omission or commission) in any condition, a negative sensitivity index or due to a high level of EEG artifacts (less than 20 remaining trials in any condition). Data of the following age groups were included: 35 adults, 18–34 years (mean: 24;8 (years;months), 26 females, 2 left-handed, 39 teens, 10–12 years (mean: 11;3), 18 females, 2 left-handed, and 30 children, 6–8 years (mean: 7;11, 18 females, 6 left-handed). Participants either received a voucher for a local toy shop and a certificate (children, 10€/hour) or payment (adults, 10€/hour). All participants gave written consent (both children, parents and adults). Participants confirmed a normal or corrected-to-normal vision, normal hearing abilities, no intake of medication with effects on the nervous system, and no history of attention-related disorders. Handedness was measured with a shortened German version of the Oldfield Handedness Inventory (Oldfield, 1971). The project was approved by the local ethics committee.

### 2.2 Design Considerations and Sample Size

In a cross-sectional study we examined RT, accuracy, pupil dilation and EEG in three age groups. We applied Bayesian statistics. Unlike frequentist p-values, which remain uniformly random under the null hypothesis regardless of sample size, BF_10_ is not random and systematically accumulates evidence in favor of the null or alternative model as the number of participants increases (Schönbrodt et al., 2017). All results that are relevant to the conclusions of this study provide clear evidence for or against the null hypothesis, indicating that the sample size is sufficient to detect significant interactions between involuntary and executive attention as well as age differences. Previous relevant studies conducting similar Go/Nogo paradigms collected data from a similar or lower number of participants (e.g. Barry et al., 2014; Jonkman et al., 2003). Similar results revealed a sample size calculation using G*Power (f=0.25, α=0.05, Power=0.8; Faul et al., 2007).

### 2.3 Stimuli, Software and Apparatus

#### Stimuli

The stimuli included four different animal sounds (cat meow, horse neigh, sheep baa and rooster crow) with a pitch transition starting 200ms after sound onset of either increasing or decreasing frequency. The pitch shift (+/- 4 semitones, transition time: 40ms) was introduced using version 3.0.0 of Audacity software. Sounds had a duration of 500ms with a 10ms rise and fall time and were equalized using root mean square normalization (of the initial 200ms).

#### Software and Apparatus

The experiment was conducted in an acoustically attenuated and electromagnetically shielded cabin. Illuminance of the cabin was held constant at a level of 48.9lx (measured with GOSSEN MAVOLUX 5032B USB, Nürnberg, Germany). Participants sat in front of the screen with a viewing distance of 55cm. The head position was supported by using a chin-rest (SR research head support, Ottawa, Ontario, Canada).

Auditory stimuli were presented via loudspeakers (IK Multimedia iLoud Micro Monitors) located at the left and the right of the screen. Sounds were presented with a loudness of 65.3dB SPL (measured with PAA3 PHONIC, Taipei, Taiwan).

Visual stimuli were presented on a 23.6” VIEWPixx EEG display (VPixx Technologies Inc.) with a resolution of 1920x1080px (23.6-in. diagonal display size) and a refresh rate of 120Hz. The experimental stimulation was presented via Psychtoolbox (Version 3.0.15, Brainard, 1997) using Octave (Linux, Version 4.0.0). Response buttons were connected with a RT-Box which provided precise response time measurements (Li et al., 2010).

### 2.4 Task and procedure

The task was introduced using the same instruction video for every participant. Participants were instructed to press a button as fast and accurate as possible (with the index finger of the dominant hand) when they hear the respective Go-sound. Go-sounds were sounds with either ascending or descending pitch change, balanced over all participants (Figure 1). Go and Nogo sounds were presented equiprobably. In each of four blocks (6min), two of four different animal sounds (pairs balanced across participants) were presented: the standard sound with an 80% probability (60/40% Go and 60/40% Nogo) and the deviant sound with a 20% probability (15/10% Go and /1510% Nogo). The order of animal-pairs (standard/deviant) presented per block was balanced across participants using latin square. Sounds were presented in pseudo-randomized order with a sound onset asynchrony (SOA, 1800/1900/2000/2100/2200ms). Each deviant was preceded by at least two standard sounds.

**Figure 1.**
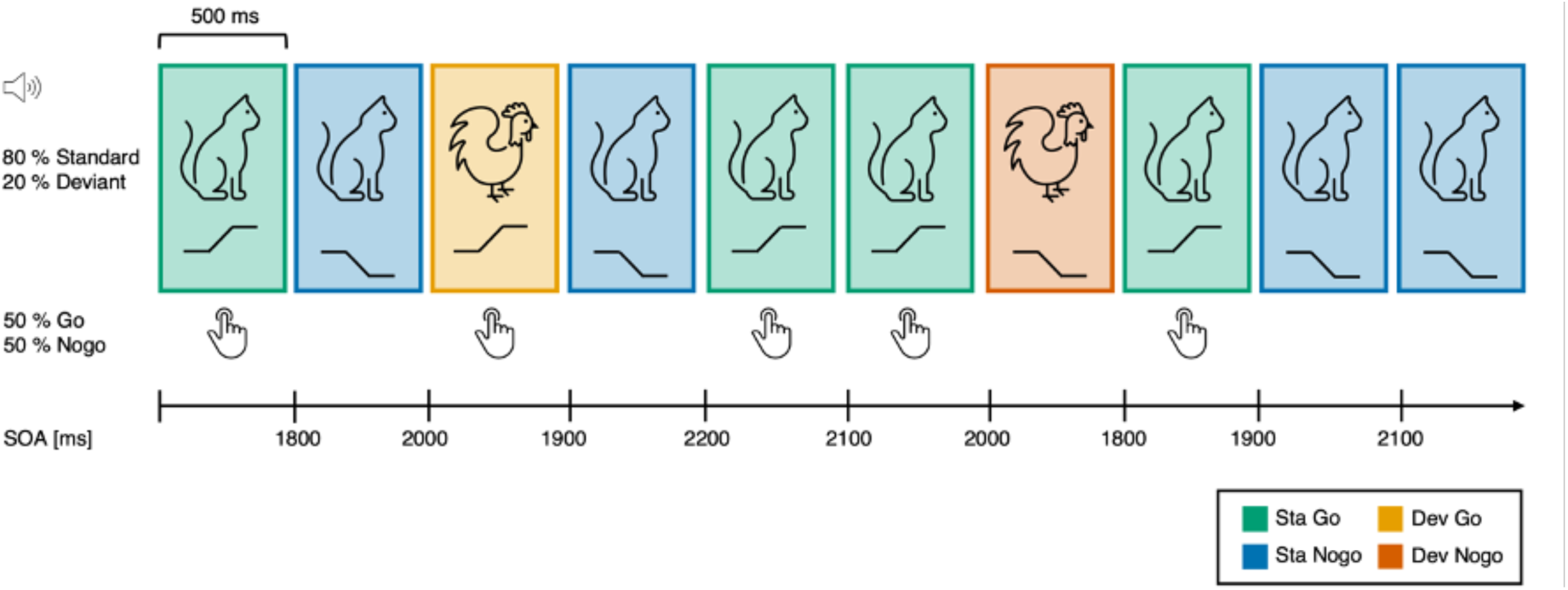
Task sound sequence. Schematic representation of the equiprobable Go/Nogo task at the beginning of a block. Animal sounds were presented with either increasing (Go) or decreasing (Nogo) pitch shift 200ms after sound onset. Different animal sounds are used as the distracting feature of deviant sounds (orange: Deviant Go, red: Deviant Nogo).

To keep motivation high, the task was embedded in a child-friendly story and combined with a game-like visual feedback at the end of each block, where food for the animal was collected with each correct answer. Illustrations were displayed at the center of a screen with a size of 35.8°x20.6° visual angle. Mean luminance of the fixation picture was 48.2cd/m^2^ on a black background screen with a mean luminance of 0.3cd/m^2^.

Each experimental block started with a five-point eye-tracker calibration and validation procedure. The visual feedback on correct button presses was presented after every 50 trials (3 times total per block), while during stimulation participants were instructed to fixate a point in the middle of the screen (the steering wheel of a tractor placed in a farm scene). This was to keep eye-movements to a minimum. Prior to each block, participants performed a short training including 12 trials (each sound-type was presented 3 times). The sounds presented in the training block were the same sounds as in the following experimental block. The training block was repeated until the participant made less than two mistakes (adults: 1, teens: 2, children: 3 trainings on average at the beginning of the experiment).

### 2.5 Data recording

The electroencephalogram (EEG) was recorded at a sampling rate of 500Hz from an ActiChamp amplifier and a 31 active electrode Braincap (Brain Products GmbH, Gilching, Germany). The electrodes were placed according to the extended 10–20 system (Fp1-Fp2- F7-F3-Fz-F4-F8-FC5-FC1-FC2-FC6-T7-C3-Cz-C4-T8-CP5-CP1-CP2-CP6-P7-P3-Pz-P4-P8- Oz) and at the left and right mastoids. Three electrodes recording the horizontal and vertical electrooculogram (EOG) were positioned to the left and right of the outer canthi of the eyes and below the left eye. The reference electrode was placed at the tip of the nose.

### 2.6 Data analyses

The first two standard trials per block and two standard trials following each deviant sound were excluded from all analyses. Only trials where participants’ responses were given correctly and in the time window from 300−1700ms after sound onset were used for behavior, pupil- and EEG data analyses.

#### EEG data processing

Pre-processing of EEG data was carried out using EEGLAB (version 2026.0.0; Delorme & Makeig, 2004). The signal was down-sampled offline to a sampling rate of 250Hz and filtered with a 48Hz low-pass filter (order 208 Hamming-windowed sinc FIR filter, transition band with 4Hz) and a 0.1Hz high-pass filter (order 4126, transition band with 0.2Hz). The continuous data were cleaned with the GEDAI-EEGLAB plugin (“generalized eigenvalue de-artifacting instrument”, version 1.5; Ros et al., 2025) with a “low-cut frequency” parameter of 0.1Hz and default setting otherwise. Data was then segmented into epochs of 1.3s including a 0.2s pre-stimulus baseline period. Epochs with peak-to-peak amplitudes exceeding 150µV at any electrode were excluded and epochs were baseline corrected (0.2s pre-stimulus). The data were re-referenced to “infinity” using the REST_cmd-EEGLAB plugin (“Reference Electrode Standardization Technique”, version 1.0; Dong et al., 2017). Mean amplitudes were computed within 40ms (Nogo-N2 effect) or 80ms (all other analyses) time windows and regions of interest (ROI) centered on the spatial and temporal peak (as derived by jack-knifing; see below) of the respective component in the ERP (P3b) or the difference wave (Nogo minus Go for Nogo- N2/P3 effects, and deviant minus standard for P3a) per group. We could not derive sufficiently precise canonical analysis time windows due to the novel experimental design but considered our approach acceptable because we based all of our conclusions on Bayesian evidence (applying default priors) provided by the data rather than frequentist statistics.

#### Residue iteration decomposition (RIDE)

We applied residue iteration decomposition (RIDE; Ouyang et al., 2015) to dissociate sensory, motor, and central ERPs components. Our analyses focused on the C-component as it reflects higher-level cognitive processes such as response inhibition (Ouyang et al., 2015). We consider the application of the RIDE decomposition as necessary to compare the ERP data between age groups due to the different variability of the RIDE C-component in our data (S3.2.1). Details are provided in the supplementary information (S3.1).

#### Jack-knifing

Component peak maximal amplitudes and corresponding latencies were analyzed by computing individual jack-knifing estimates separately for each group (Kiesel et al., 2008). Latencies were extracted from these waveforms at the time point where the amplitude reached the peak amplitude. Individual latencies were then calculated from the subsample latencies as suggested by (Smulders, 2010) to account for the reduced variance in the jack-knifing estimates (Smulders, 2010; Equation 1). Those individual latencies were then subjected to Bayesian statistics.

### 2.7 Statistical analyses

#### Sensitivity analysis

The d-prime sensitivity index (d’) and the response criterion (c) was computed per participant and sound type from the hit rate (proportion of correct Go responses) and false alarm rate (proportion of incorrect Nogo responses). We applied log-linear correction of individual hit and false alarm rates (Stanislaw & Todorov, 1999).

#### Bayesian analyses

Statistical analyses on behavioral, PDR (S2) and ERP data were performed using JASP (version 0.96; JASP Team, 2023). Separate repeated measures Bayesian ANOVAs were calculated for each dependent variable: RT, RT variability (S1.1), sensitivity index d’, criterion c, PDR amplitude and latencies (S2.2), as well as ERP S- or C-components (RIDE decomposition) and classic ERP components (S4.1) Nogo-N2 effect, Nogo-P3 effect, P3a and P3b amplitudes and latencies. The ANOVAs include sound type (levels: standard, deviant) and target (levels: Go, Nogo) as within-subjects factors (where applicable) and age group as between-subjects factor (children, teens, adults). The JASP default fixed and random effects priors were used, defined as, respectively, *r*=0.5 and *r*=1. We report Bayes factor (BF_10_), where larger values suggest that more evidence is provided by the data for a model reflecting the alternative hypothesis (lower for model reflecting the null hypothesis). BF_10_ was interpreted following Lee & Wagenmakers (2013), where moderate evidence in favor of the alternative (or null) hypothesis is considered, if BF_10_ is larger than 3 (or lower than 0.33), or strong evidence if BF_10_ is larger than 10 (lower than 0.1). BF_10_ between 0.33 and 3 are considered as weak or “anecdotal” evidence. In addition, inclusion Bayes factors based on matched models were calculated by comparing the models containing a main or interaction effect to the equivalent models stripped of the effect (BF_incl_; Mathôt, 2017). Main effects and interactions were analyzed using Bayesian independent samples (for between-subjects factor) or paired samples *t*-tests (for within-subject factors), where the null hypothesis corresponded to a standardized effect size δ=0, while the alternative hypothesis was defined as a Cauchy prior distribution centered around 0 with a scaling factor of *r*=0.707.

## 3 Results

### 3.1 Behavior

**Response times (RT)** to Go stimuli decreased with age. All age groups showed a distraction effect: slower response times for deviants compared to standards (Table 1, Figure 2)This distraction effect decreased with age (Ch sta vs. dev: 838 vs 942ms; Te: 744 vs 853ms; Ad 622 vs 678ms; relative to sound onset).

**Figure 2.**
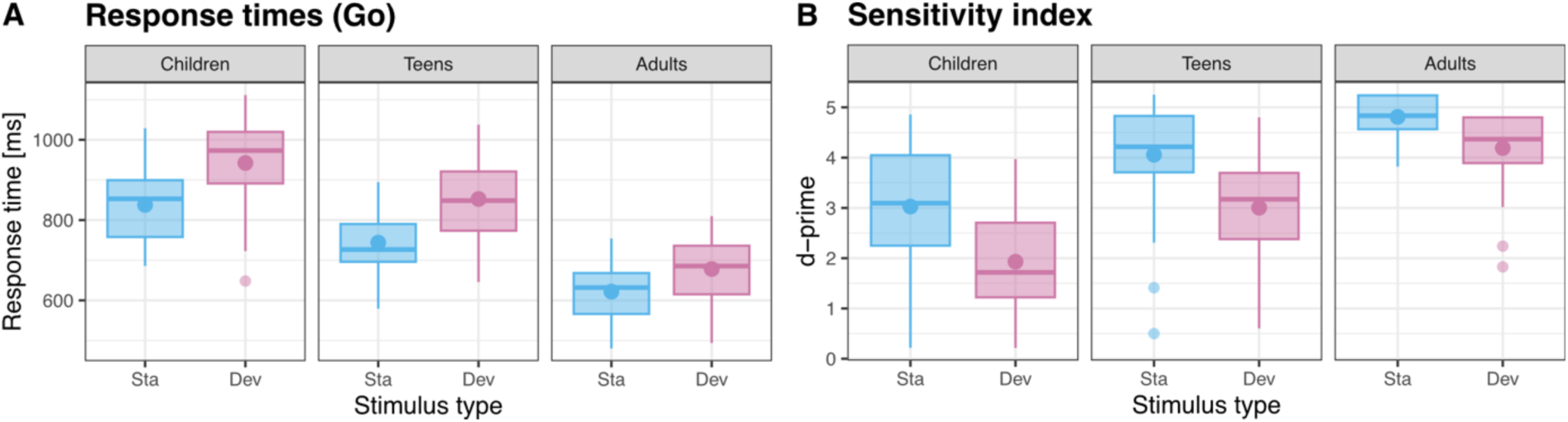
Response times for standard and deviant Go sounds (Panel A) and d’ sensitivity index for standard and deviant sounds (Panel B), in children, teens and adults.

**Table 1.** Behavioral performance per age group and sound type.

| Group | Sound type | RT (ms) | Go hit rate | Nogo CR rate | d' | c |
| --- | --- | --- | --- | --- | --- | --- |
| Children | Standards | 838 | 0.895 | 0.892 | 3.03 | -0.02 |
|  | Deviants | 942 | 0.778 | 0.798 | 1.93 | 0.05 |
| Teens | Standards | 744 | 0.970 | 0.957 | 4.06 | -0.05 |
|  | Deviants | 853 | 0.928 | 0.888 | 3.00 | -0.09 |
| Adults | Standards | 622 | 0.998 | 0.989 | 4.81 | -0.13 |
|  | Deviants | 678 | 0.980 | 0.969 | 4.19 | -0.06 |

The Bayesian ANOVA preferred the model including the sound type and age group main effects and the sound type by age group interaction effect (BF_10_=8.97×10^38^). The data provided strong evidence for a distraction effect in all groups (Ch: BF_10_=1.12×10^5^; Te: BF_10_=1.33×10^9^; Ad: BF_10_=5.08×10^7^). The data provided moderate evidence against a difference in the distraction effect between children and teens (BF_10_=0.254) and strong evidence for a difference in the distraction effect between adults and children (BF_10_=12.4) or teens (BF_10_=136). The data provided strong evidence for RT differences between all age groups (Ch >Te: BF_10_=454; Ch >Ad: BF_10_=1.85×10^14^; Te >Ad: BF_10_=1.64×10^9^).

**The sensitivity index d’** increased with age. All age groups showed a distraction effect: lower d’ for deviants compared to standards. This distraction effect decreased with age (Table 1, Figure 2). The Bayesian ANOVA preferred the model including the sound type and age group main effects and the sound type by age group interaction effect (BF_10_=2.74×10^32^). The data provided strong evidence for a distraction effect in all age groups (Ch: BF_10_=2.3×10^8^; Te: BF_10_=2.46×10^8^; Ad: BF_10_=2487). The data provided moderate evidence against a difference in the distraction effect between children and teens (BF_10_=0.257) and moderate evidence for a difference in the distraction effect between adults and children (BF_10_=8.69) or teens (BF_10_=4.26). The data provided strong evidence for d’ differences between all age groups (Ch vs Te: BF_10_=283; Ch vs Ad: BF_10_=5.54×10^10^; Te vs Ad: BF_10_=2.78×10^4^).

Importantly, **the criterion c** did neither systematically change with age nor sound type. The Bayesian ANOVA preferred the null model. The data provided anecdotal evidence against a main effect of age group (BF_incl_=0.373) as well as moderate evidence against a main effect of sound type (BF_incl_=0.209) and a sound type by age group interaction effect (BF_incl_=0.183). In other words, the tendency towards responding was not enhanced for deviants compared to standards in any age group.

In sum, with age, RTs, RT variability (S1.1) and false alarm rate (S1.1) decreased; while sensitivity and hit rate (S1.1) increased. RTs were higher and sensitivity lower for deviants than standards, in all age groups. Distraction effects on RT and sensitivity were smaller in adults. The response criterion was not affected by sound type or age group.

### 3.2 ERP

Figure 3 displays the ERPs in each condition for each age group and the decomposition into the sensory (S-component, locked to stimulus onset), the motor processing (R-component, locked to button press) and the central (C-component, variable latency) components.

**Figure 3.**
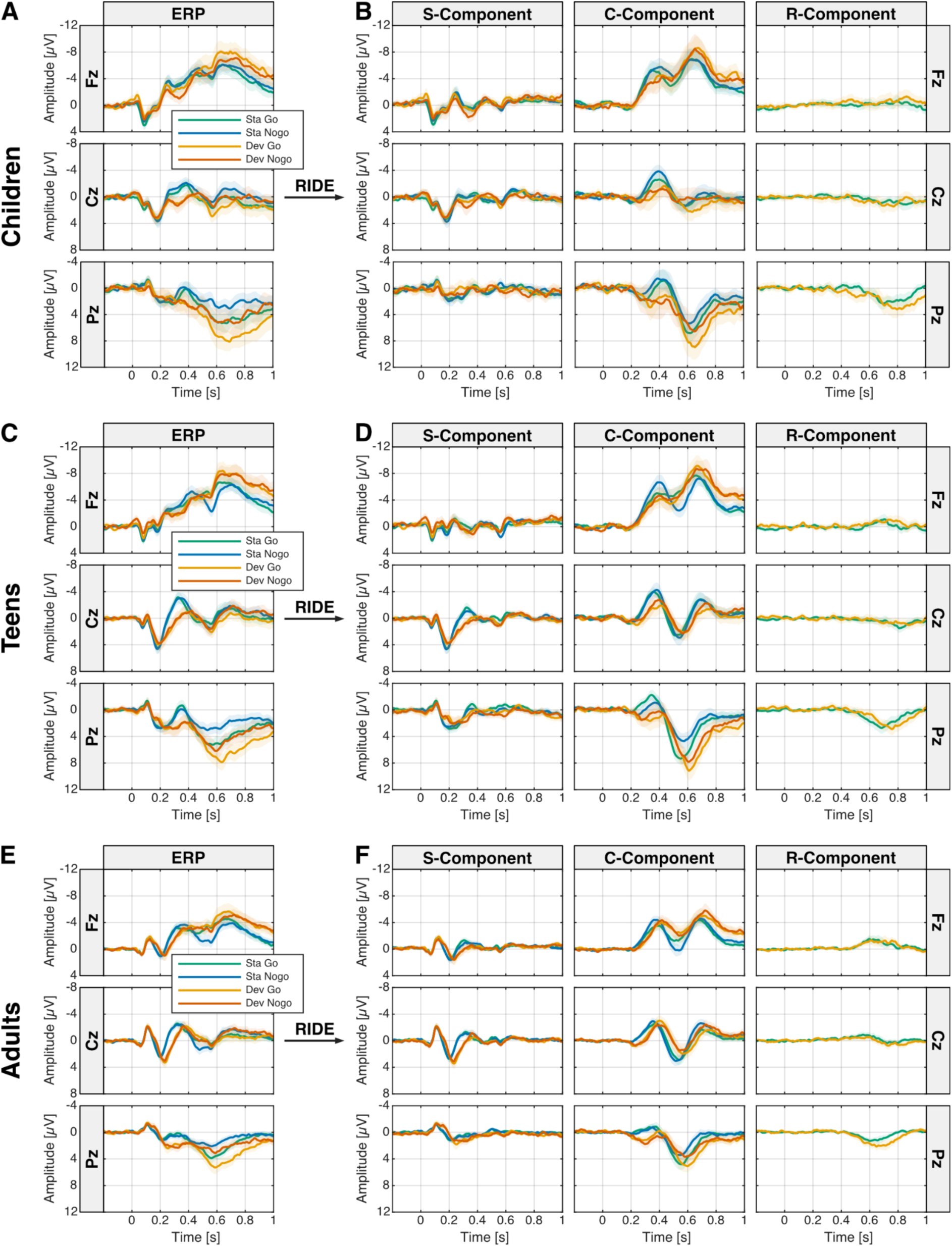
ERP response and RIDE decomposition. Left: Grand averages of the original ERP response for children (A), teens (C) and adults (E) for each condition. Right: Waveforms of the S-component (stimulus locked), C-component (variable latency) and R-component (response locked) after RIDE decomposition for children (B), teens (D) and adults (F). Shaded areas (in all figures) indicate 95% confidence intervals.

#### RIDE decomposition

The S-component showed the typical expected sensory waves as in the ERPs: the fronto-central P1-N1-P2 waves in adults, the fronto-central P1-P2 wave in the 6–8-year olds, and a pattern in between in 10-12-year-olds (Figure 3, (Bruneau et al., 2015; Ponton et al., 2000; Wunderlich & Cone-Wesson, 2006)). The central P3a was best captured by the S-component (dev minus std difference) and was observed for both Go and Nogo stimuli, in all age groups (S3.2.2&4.1.1). In the R-component, a long late parietal positivity was apparent in all groups, which likely reflects both the motor response and the somatosensory feedback of the button press (Ouyang et al., 2013).

To identify higher-level cognitive processes, we analyzed the C-component after RIDE decomposition (Ouyang et al., 2015). In all groups, the C-component showed a pattern consisting of a frontal negative wave peaking around 400ms and a parietal positive wave peaking between 600 and 700ms, resembling the N2 and P3 which are typically observed in Go/Nogo paradigms. We computed the Nogo–Go difference wave at Fz electrode location showing frontal Nogo-N2 effect and Nogo-P3 effect in the response to standard sounds (Figure 4&5). Note, that the stimulus feature indicating a Go or NoGo response was presented 200ms after sound onset.

**Figure 4.**
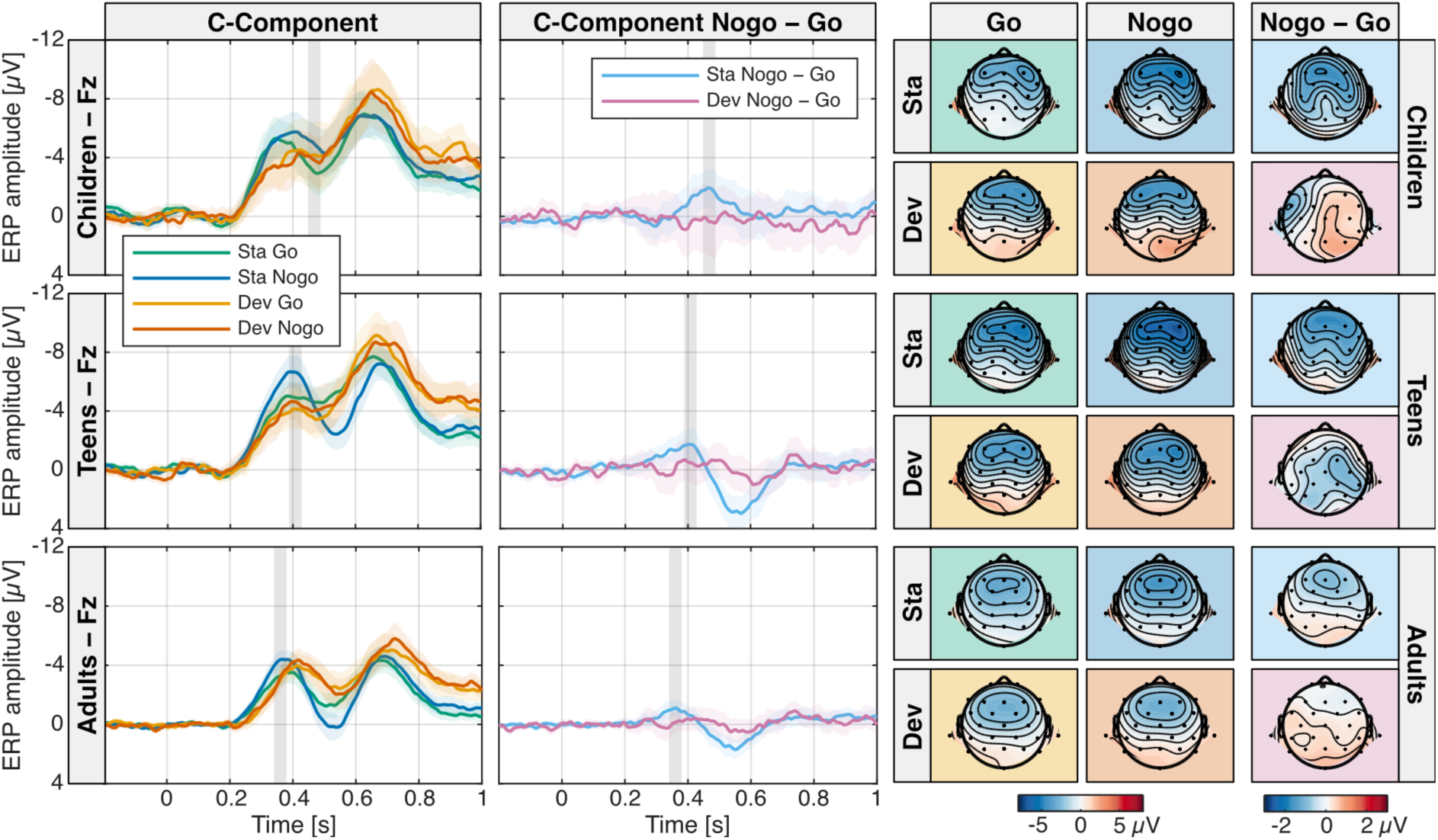
Nogo-N2 effect in the RIDE C-component. A frontal Nogo-N2 effect was observed in the Nogo minus Go difference wave for standards but not for deviants in all age groups. The latency of the Nogo-N2 effect decreased with age. Left: RIDE C-component ERP waves per condition and age group, Middle: Corresponding Nogo minus Go difference waves. Grey bars indicate the analysis time window. Right: RIDE C-component ERP topographies in the Nogo-N2 effect time window per condition and age group and corresponding Nogo minus Go difference topographies.

#### Nogo-N2 effect

We observed more negative C-component amplitudes around 350-450ms over frontal areas in response to Nogo stimuli compared to Go stimuli for Standards but not for Deviants. We consider this difference potential which is best visible in the Nogo minus Go difference wave (Figure 4) as the Nogo-N2 effect. The peak latency of the Nogo-N2 effect for standards was 468ms for children, 408ms for teens, and 360ms for adults with peak amplitudes at Fz electrode location in all age groups (Figure 4). The amplitudes of the Nogo-N2 effect for standards were similar between age groups. We could not identify a corresponding Nogo-N2 effect in deviant trials.

##### Latency

The latencies were therefore analyzed only in standard trials. The Bayesian ANOVA on the standard Nogo-N2 effect latency preferred the model including the group main effect (BF_10_=2.45×10^7^). There was strong evidence for all Nogo-N2 effect latency differences between age groups (Ch > Te: BF_10_=26.9, Ch > Ad: BF_10_=4.35×10^13^, Te > Ad: BF_10_=35.0).

##### Amplitude

As no clear peak for a Nogo-N2 effect in deviants could be identified, we computed the mean amplitudes in (40ms duration) time windows centered on the peak of the Nogo-N2 effect in standards per group. The Bayesian ANOVA on amplitudes preferred the model including the sound type, target and age group main effects and the sound type by target interaction effect (BF_10_=1008). The data provided moderate evidence against a target by age group (BF_incl_=0.106) and a sound type by target by age group interaction effect (BF_incl_=0.232). There was strong evidence for a Nogo-N2 effect in standards (Nogo > Go: BF_10_=1.74×10^4^), but moderate evidence against a Nogo-N2 effect in deviants (BF_10_=0.114).

In sum, we observed larger amplitudes in Nogo than Go trials for standards, but not for deviants, with similar amplitude in all age groups at latencies decreasing with age.

#### Nogo-P3 effect

We observed more positive C-component amplitudes around 500-600ms over frontal areas in response to Nogo stimuli compared to Go stimuli for standards but not for deviants in teens and adults. We consider this difference potential which is best visible in the Nogo minus Go difference wave (Figure 5) as the Nogo-P3 effect. The Nogo-P3 effect peak latency for standards was 568ms for teens, and 552ms for adults with peak amplitudes at Fz electrode location in both age groups (Figure 5). We could not identify a corresponding Nogo-P3 effect in deviants. In children, we could not identify a Nogo-P3 effect, neither in standard nor in deviant trials.

**Figure 5.**
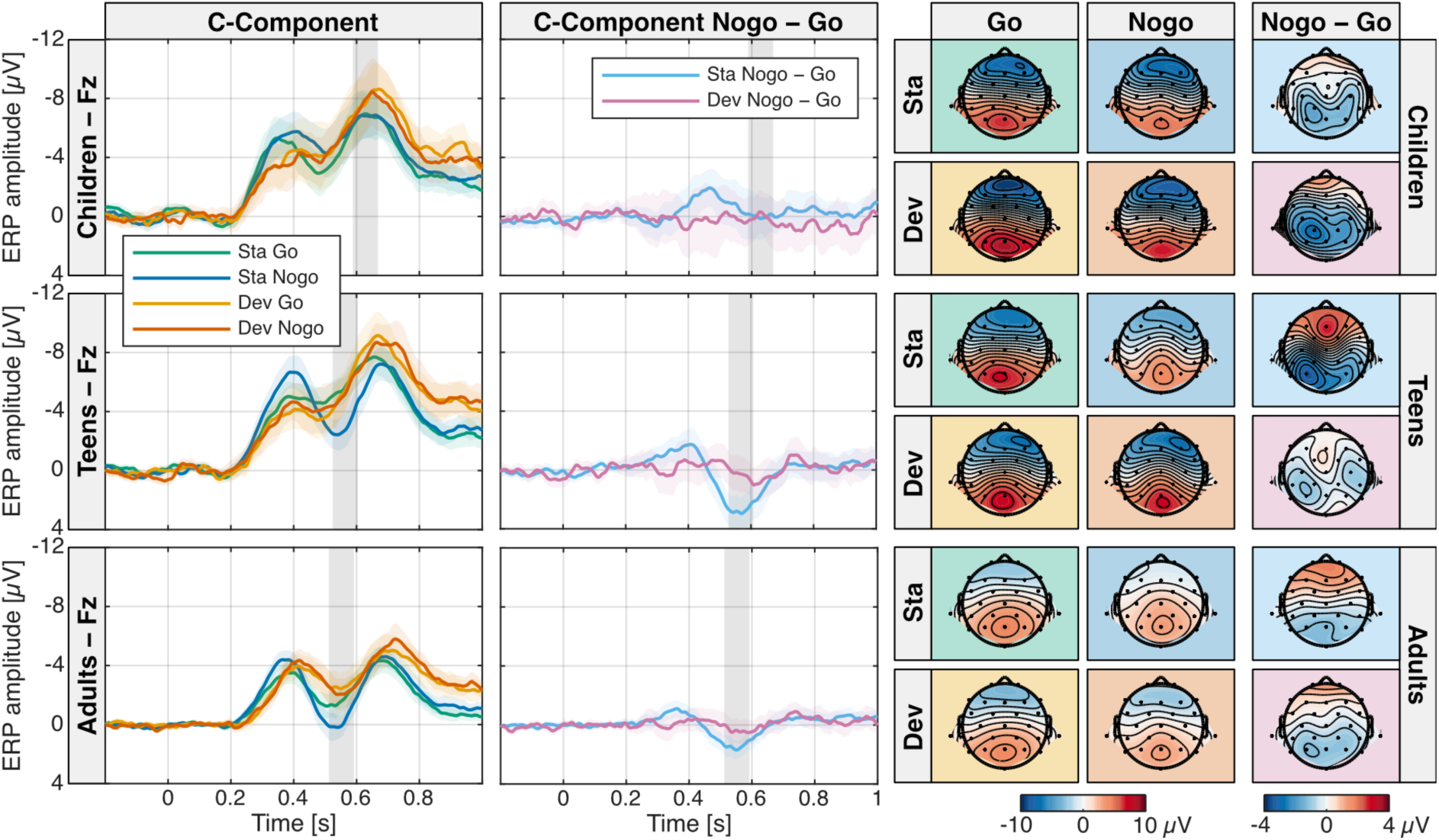
Nogo-P3 effect in the RIDE C-component. A frontal Nogo-P3 effect was observed in the Nogo minus Go difference wave for standards but not for deviants in teens and adults but not in children. Left: RIDE C-component ERP waves per condition and age group, Middle: Corresponding Nogo minus Go difference waves. Grey bars indicate the analysis time window. Right: RIDE C-component ERP topographies in the Nogo-P3 effect time window per condition and age group and corresponding Nogo minus Go difference topographies.

##### Latency

The data provided moderate evidence against a latency difference of the Nogo-P3 effect between teens and adults (BF_10_=0.260).

##### Amplitude

As no clear peak for a Nogo-P3 effect in deviants could be identified, we computed the mean amplitudes in (80ms duration) time windows centered on the peak of the Nogo-P3 effect in standards in teens and adults, respectively, and the most positive peak in the corresponding time range at 628ms in children. The Bayesian ANOVA on the amplitudes preferred the model including the sound type, target and age group main effects and the sound type by target and target by age group interaction effects (BF_10_=2.13×10^13^). The data provided moderate evidence against a sound type by age group (BF_incl_=0.096) and anecdotal evidence for a sound type by target by age group interaction effect (BF_incl_=2.18). There was strong evidence for a Nogo-P3 effect in standards (Nogo > Go: BF_10_=3749), but moderate evidence against a Nogo-P3 effect in deviants (BF_10_=0.168). There was moderate evidence against a Nogo-P3 effect in children (Nogo > Go: BF_10_=0.195), but strong evidence for a Nogo-P3 effect in teens (BF_10_=83.2) and adults (BF_10_=10.7).

In sum, we observed a Nogo-P3 effect for standards in teens and adults, but not in children nor for deviants in any age group.

To verify that all reported effects were not introduced by the RIDE decomposition, we performed all analyses also on the individual and grand-average ERPs. Effect sizes were reduced, but all results relevant for the conclusion were successfully replicated (except the latency effects for Nogo-N2 effects; presumably due to the considerably higher noise level of the ERPs). As the ERP analyses do not contribute to (or contradict) our conclusions, we do report them in the supplementary materials (S4).

## 4 Discussion

The present study examined the developmental pathway of the impact of attentional distraction on executive attention during a combined auditory Go/Nogo oddball paradigm in children, adolescents and adults. Effects of involuntary orienting of attention evoked by deviant sounds were observed at behavioral and neural levels. Distraction effects decreased with age. In the EEG, a frontal Nogo-N2 effect was observed in all age groups during standard trials but not during deviant trials. Moreover, a frontal Nogo-P3 effect was observed in adolescents and adults, during standard trials only, but was completely absent in children. Finally, remarkable age differences in the processing of the standard and deviant sounds (P3b) and pupil dilation responses to deviant and target sounds are presented and discussed in the supplementary material.

### 4.1 Age effects on involuntary and executive attention

Our findings at the behavioral (Figure 2) and the pupillometry levels (Fig S2) confirmed the expected distraction effect in all groups: slower and less accurate responses (RT, d’) to deviant compared to standard sounds. This indicates involuntary orienting of attention towards unexpected and task-irrelevant deviants, resulting in impaired task-performance in all age groups (Escera et al., 1998; Schröger & Wolff, 1998; Wetzel et al., 2006). Distraction effects were largest in children and teens compared to adults. These results are in line with previous findings indicating immature attention in 6−8-year-olds (Hoyer et al., 2021; Wetzel et al., 2019), and suggest on-going maturation of attention processes during adolescence (Booth et al., 2003; Hoyer et al., 2021; Petton et al., 2019; Thillay et al., 2015). In the EEG, a clear fronto-central P3a component was observed in all groups demonstrating that deviants involuntarily captured attention and increased arousal (Escera et al., 2000; Masson & Bidet-Caulet, 2019; Fig S3&S5).

We assessed executive attention processes typically attributed to the Go/Nogo task including response inhibition and target processing on the behavioral and pupillometry levels. In traditional tasks with unequal probability of Go/Nogo trials, false alarms (FA) are the main behavioral measure for response inhibition. As expected, we observed also in our equiprobable Go/Nogo task a decreasing FA rate with age (Fig S1), indicating an ongoing maturation of response inhibition after the age of 12. These findings are consistent with an improvement of response inhibition from childhood through adolescence until young adulthood (Durston et al., 2002; Humphrey & Dumontheil, 2016; Johnstone et al., 2005; Motes et al., 2018). Importantly, the sensitivity analysis revealed, that the response tendency was not increased for deviants compared to standards in any age group. Thus, an interaction of orienting and executive attention cannot be explained by a response bias in favor of deviant sounds.

### 4.2 Age-related modulation of the interaction between involuntary attention and executive attention

A Nogo-N2 effect and a Nogo-P3-effect, that have been associated with inhibition processes, were observed. There was very strong evidence for a Nogo-N2 effect during standard trials in all age groups, indicating successful response inhibition (Cheng et al., 2019). Importantly, a Nogo-N2 effect was absent during deviant trials in all age groups. The Bayesian statistics confirmed that the lack of the Nogo-N2 effect in deviant trials is not due to a lack of power (BF_10_=0.114). This finding suggests that deviant-related attention processes affect response inhibition in the time range of the frontal N2. It can be assumed that available cognitive resources are initially directed toward the processing of the unexpected sensory input (deviant) at the cost of inhibition processes at the brain level. This finding confirms our hypothesis of an interaction between involuntary and executive attention. However, our data indicate evidence against age differences in this interaction throughout school age.

In contrast to previous studies using unequal probabilities of Go and Nogo trials (Hoyniak, 2017, Jonkman, 2006; Johnstone, 2005), the amplitude of the Nogo-N2 effect did not decrease with age in our study. This could be caused by the equiprobable design of the present study that initiates no strong response bias. In line with previous studies (Hoyniak, 2017), the latency of the Nogo-N2 effect in standard trials clearly decreased with age by about 100ms from children to adults. This can be attributed to immature brain structures involved in response inhibition. In particular, the myelinization of axons lasts into adulthood and increases processing speed (Giedd et al., 2015) and presumably response inhibition capacities.

A frontal Nogo-P3 effect was observed in teens and adults only. The absence of the Nogo-P3 effect in children was confirmed by Bayesian statistics (BF_10_=0.195). While studies using unequal Go/Nogo probabilities revealed inconsistent findings regarding the development and topography of the Nogo-P3 effect in children (Hoyniak, 2017), the few equiprobable Go/Nogo studies with children consistently reported a frontal Nogo-P3 effect in 8−13- and 9−11-year-olds (Barry et al., 2014; Barry et al, 2018). The present results further specify the developmental pathways of the underlying inhibition mechanisms by showing immature inhibition capabilities in 6−8-year-old children. For older children, the present findings confirm the advanced development of response inhibition in early adolescence in Go/Nogo tasks that elicit no strong response tendency.

Importantly, the Nogo-P3 effect in the older groups was confirmed in standard trials only. This indicates that deviant-related attention processes affect response inhibition also in the time range of the Nogo-P3 effect. Both, Nogo-N2 and Nogo-P3 effects have been assumed to reflect response inhibition (Cheng, 2019). In contrast to the Nogo-N2 effect, the subsequent Nogo-P3 was found larger when the response included a motor response compared to counting of targets (Bruin Wijers; 2002; Smith, 2008). The authors suggested that the inhibition of a manual response requires a higher level of inhibition than counting and that the Nogo-P3 effect is a marker of motor inhibition. Different mechanisms underlying the Nogo-N2 and Nogo- P3 effect are also evidenced by the distinct developmental patterns observed in young children in the present study. In 6−8-year-olds, while the amplitude of the Nogo-N2 effect was similar to that of adults, the Nogo-P3 effect was completely absent. Moreover, and in contrast to the Nogo-N2 effect, the latencies of the Nogo-P3 effect were similar between teen and adult age groups. These findings support the assumption that different inhibition mechanisms underlie the Nogo-N2 and Nogo-P3 effects and suggest that these mechanisms follow different developmental trajectories.

The distinct maturation of the discussed inhibition mechanisms, particularly between children and adolescents, is not directly reflected in task-performance as distraction effects (RTs as well as task performance as reflected in d’) were similar between children and adolescents but reduced in adults. This suggests the contribution of further relevant mechanisms involved in the inhibition of task-irrelevant deviants and the facilitation of task-relevant targets.

### 4.3 Conclusions

The distinct mechanisms of response inhibition underlying the frontal Nogo-N2 and Nogo-P3 effects are affected by deviant processing. This demonstrates the interaction of involuntary orienting and executive attention. Unexpectedly, even mature inhibition mechanisms in adults remain susceptible to involuntary attentional allocation to unexpected stimuli. Significant age differences have been observed for different aspects of inhibition. While the Nogo-N2 effect underlying inhibition mechanisms are delayed in 6−8-year-old children but fully functioning at the adult level during the present task; the mechanisms underlying the Nogo-P3 effect, presumably contributing to motor inhibition, develop rapidly between 6−8 and 10−12 years. The present results demonstrate distinct developmental trajectories of the inhibition mechanisms underlying Nogo-N2 and Nogo-P3 effects throughout childhood and adolescence. Results offer valuable insights into how executive and involuntary attention interact during development. Additionally, our study introduces a promising auditory paradigm for investigating this interaction of attention processes in developmental populations.

## Data Availability Statement

Preprocessed data and statistical scripts will be made available in an appropriate repository.

## Funding

This work was supported by the DFG [WE5026/4-1], the French National Research Agency [ANR-19-FRAL-0007-01] and the Leibniz Association [P58/2017].

## Acknowledgements

We sincerely thank all participants – adults, children and their parents – whose contributions were essential to this study. We are grateful to Dunja Kunke, Gabriele Schöps and the student assistants for their efforts supporting data acquisition.

## Author Contribution

**Ursula Schöllkopf:** Conceptualization, Data curation, Formal analysis, Investigation, Methodology, Software, Visualization, Writing – original draft, Writing – review & editing. **Andreas Widmann:** Conceptualization, Data Curation, Formal analysis, Methodology, Software, Visualization, Writing – original draft, Writing – review & editing. **Aurélie Bidet- Caulet:** Conceptualization, Funding Acquisition, Supervision, Writing – original draft, Writing – review & editing. **Nicole Wetzel:** Conceptualization, Funding Acquisition, Project administration, Resources, Supervision, Writing – original draft, Writing – review & editing.

## Conflict of Interest

The authors declare that they have no competing financial interests or personal relationships that could have appeared to influence the work reported in this paper.

## Notes

### Competing Interest Statement

The authors have declared no competing interest.

