## Supplementary Material for "Interaction of involuntary and executive attention during development"

**Interaction of involuntary and executive attention during development – Supplementary materials**

^c^Dynamics of Communication and Auditory Processes Group, Aix Marseille Univ, INSERM, INS, Inst Neurosci Syst, 27 Boulevard Jean Moulin, 13005 Marseille, France

^d^University of Applied Sciences Magdeburg-Stendal, Osterburgerstraße 25, 39576 Stendal, Germany

^e^Center for Behavioral Brain Sciences, Otto-von-Guericke-Universität Magdeburg, Germany

**Overview**

The supplemental material reports methods, results and discussions that have not been reported in the main manuscript:

1 Results on performance: RT variability, hit and false alarm rates
2 Pupil dilation responses: methods, results, discussion
3 ERPs – RIDE decomposition: methods, results and discussion of the S-component P3a and C-component P3b
4 ERPs – non-RIDE: results for the S-component P3a, C-components Nogo-N2-effect, Nogo-P3 effect, C-component P3b

### 1 Performance

**1.1 Results**

**RT variability** decreased with age (Ch: *SD*=224 ms; Te: *SD*=190 ms; Ad: *SD*=114 ms) and was higher for deviants compared to standards (sta: *SD*=191 ms; dev: *SD*=158 ms). The Bayesian ANOVA preferred the model including the sound type and age group main effects (BF_10_=5.95×10^21^). The data provided anecdotal evidence against a sound type by group interaction effect (BF_10_=0.442). The data provided moderate to strong evidence for differences in RT variability between all age groups (Ch vs Te: BF_10_=7.35; Ch vs Ad: BF_10_=2.45×10^12^; Te vs Ad: BF_10_=6.72×10^8^).

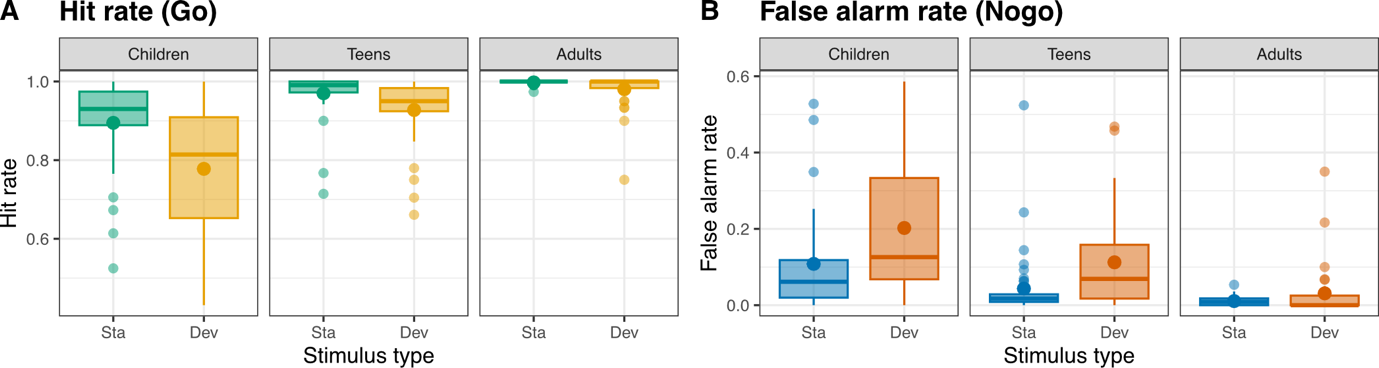

**Figure S1:** Hit rates (Go; Panel A) and false alarm rates (Nogo; Panel B) for standard and deviant Go sounds in children, teens and adults.

**The hit rate** for Go sounds increased with age and was higher for standards compared to deviants (distraction effect). The distraction effect decreased with age. The Bayesian ANOVA preferred the model including the sound type and age group main effects and their interaction (BF_10_=5.48×10^15^). The data provided strong evidence for differences in the hit rate between all age groups (BF_incl_=2.02×10^5^; Ch vs Te: BF_10_=951; Ch vs Ad: BF_10_=1.22×10^6^; Te vs Ad: BF_10_=30.3) and between standards and deviants (BF_incl_=7.60×10^7^). The distraction effect differed between all age groups (BF_incl_=359; Ch vs Te: BF_10_=9.85; Ch vs Ad: BF_10_=97.3; Te vs Ad: BF_10_=1.70). The data provided strong evidence for a distraction effect in children (BF_10_=233) and teens (BF_10_=1580) and only anecdotal evidence in adults (BF_10_=1.36).

**The false alarm rate** for Nogo sounds decreased with age and was higher for deviants compared to standards (distraction effect). The distraction effect decreased with age. The Bayesian ANOVA preferred the model including the sound type and age group main effects and their interaction (BF_10_=9.92×10^9^). The data provided strong evidence for differences in the false alarm rate between all age groups (BF_incl_=5912; Ch vs Te: BF_10_=4.61; Ch vs Ad: BF_10_=7492; Te vs Ad: BF_10_=22.5) and between standards and deviants (BF_incl_=3.98×10^5^). The distraction effect did not differ between children and teens but between children and adults and teens and adults (BF_incl_=4.07; Ch vs Te: BF_10_=0.366; Ch vs Ad: BF_10_=16.6; Te vs Ad: BF_10_=2.50). The data provided strong evidence for a distraction effect in children (BF_10_=233) and teens (BF_10_=131) and anecdotal evidence against a distraction effect in adults (BF_10_=0.773).

#### 2 Pupil dilation responses

**2.1 Methods**

#### Pupil data recording and processing. The pupil diameter of both eyes was recorded at a sampling rate of 500 Hz with an infrared EyeLink Portable Duo eye-tracker (SR Research Ltd., Mississauga, Ontario, Canada) in head-stabilized tracking mode.

Pupil analysis was implemented with MATLAB software. Eye tracker pupil diameter digital counts were calibrated using the method suggested by Steinhauer et al. (2022) and converted to mm. Blinks and enclosing saccades were marked by the blink events provided by the eye-tracker. Partial blinks (not reported by the eye-tracker) were detected from the smoothed velocity times series as pupil diameter changes exceeding 20 mm/s including a 50 ms pre-blink and a 100 mspost-blink interval using an additional custom function (Merritt et al., 1994). The data from both eyes were averaged using the dynamic offset algorithm suggested by Kret & Sjak-Shie (2019). Blinks and partial blinks (and intervals with signal loss) shorter than 1 s were interpolated using piecewise cubic Hermite polynomial interpolation. Missing data segments longer than 1 s were discarded from the data. Data was further segmented into data epochs of 2 s duration (including a 0.2 s pre-stimulus baseline). Epochs were baseline corrected by subtracting the mean amplitude of the baseline period from each epoch (Murphy et al., 2014; Widmann et al., 2018). Individual average pupil dilation responses (PDRs) were computed and peak amplitude and latency were extracted per participant and sound type (standard-Go, standard-Nogo, deviant-Go, deviant-Nogo) within a time window between 0.3-1.8 s after stimulus-onset which were then subjected to statistical analysis.‬‬‬‬‬‬‬‬‬‬‬‬‬‬‬‬‬‬‬‬‬‬‬‬‬‬‬‬‬‬‬‬‬‬‬‬‬‬‬‬‬‬‬‬‬‬‬‬‬‬‬‬‬‬‬‬

**2.2 Results**

###### **Amplitude.** PDR peak amplitudes were higher for Go compared to Nogo trials and for deviants compared to standards, for all groups (Figure S2 Panel D). The Bayesian ANOVA favored the model including sound type and target main effects (BF_10_=4.76×10^14^). The data provided strong evidence for the sound type effect (BF_incl_=6.64×10^13^) and the target effect (BF_incl_=9.83×10^24^). The data provided moderate evidence against a target by group interaction effect (BF_incl_=0.210) and strong evidence against a sound type by group interaction effect (BF_incl_=0.064) and against a sound type by target by age group interaction effect (BF_incl_=0.014).

###### **Latency.** PDR peak latencies were longer for deviants compared to standards and for Go compared to Nogo trials for all groups. This target effect was smaller in adults than in younger groups (Figure S2 Panel E). Peak latencies were decreasing with age. The Bayesian ANOVA favored the model including the sound type, target and group main effects and the target by group interaction effect (BF_10_=2.07×10^34^). The data provided strong evidence for the sound type effect (BF_incl_=2.67×10^5^), the target effect (BF_incl_=1.07×10^23^) and the group effect (BF_incl_=5.95×10^4^), and moderate evidence for the target by group interaction (BF_incl_=18.6). There was strong evidence for the target effect for each group (adults: BF_10_=8.29×10^4^; teens: BF_10_=6.85×10^9^; children: BF_10_=1.45×10^7^). There was strong evidence for a smaller Go-Nogo difference (lat_Go_–lat_Nogo_) in adults than in teens (BF_10_=49.0) and children (BF_10_=51.2) (Figure S2 Panel F). The data further suggested that the Go-Nogo difference is similar rather than different between teens and children (BF_10_=0.213). There was strong evidence for a shorter peak latency in adults than children (BF_10_=2.35×10^4^) and teens (BF_10_=786), but weak evidence for a difference between teens and children (BF_10_=0.571).

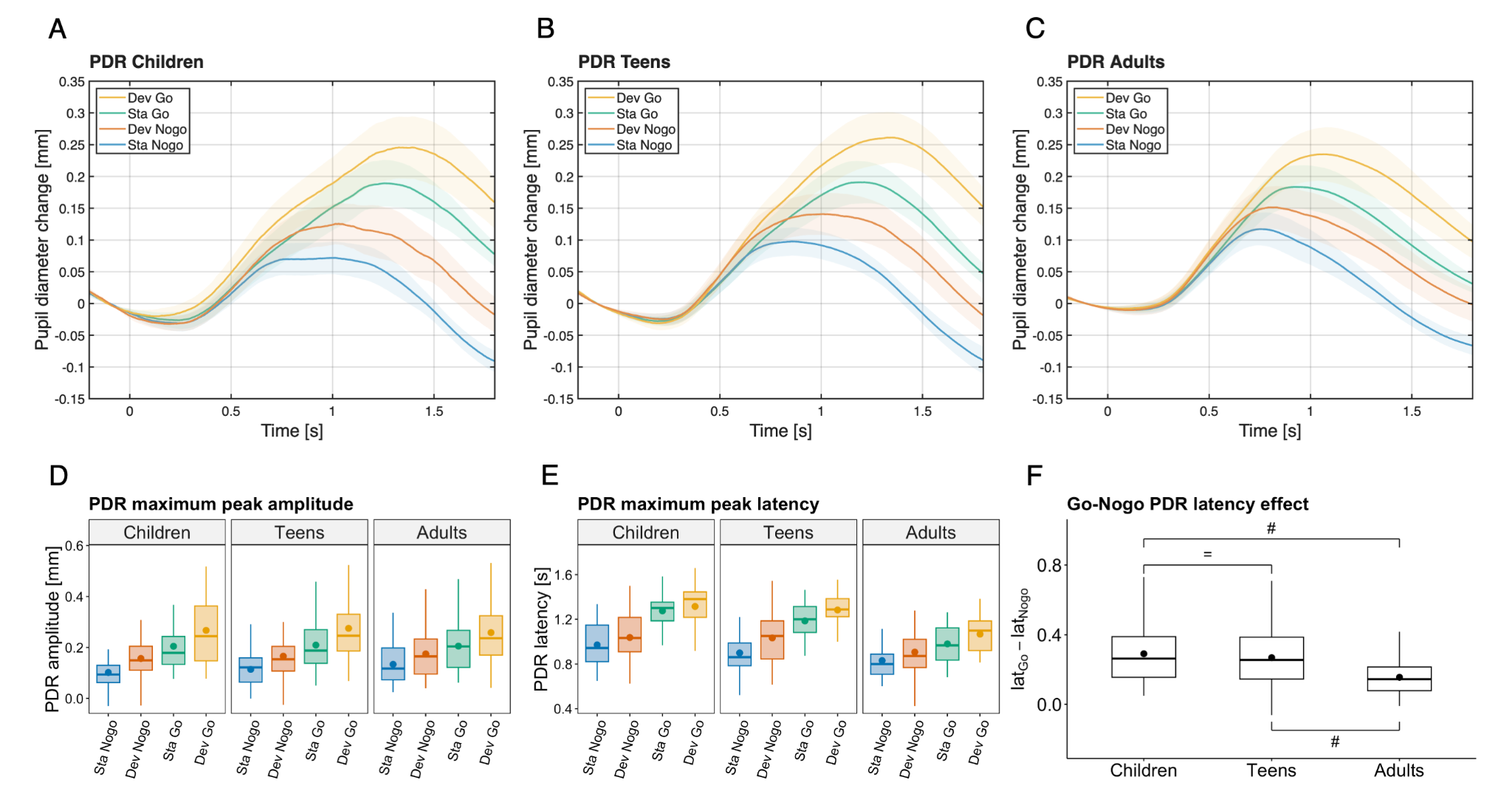

**Figure S2: Pupil dilation response (PDR).** Top: Grand average PDRs for children (A), teens (B) and adults (C) for each condition (standard Go: green, standard Nogo: blue, deviant Go: yellow, deviant Nogo: red). Bottom: PDR peak amplitude (D) and latency (E) for each condition and group. Go - Nogo latency differences (F) for each group, with evidence in favor of H1 (#) and evidence in favor of H0 (=). Boxplots display the median (central line) and mean (filled dots).

##### **2.3 Discussion**

###### 2.3.1 Age effects on involuntary attention

Both PDR amplitude and latency were larger in response to deviant than standard sounds, and thus sensitive to the unexpected sound-change in all groups. Increased amplitudes in response to oddball sounds relative to standard sounds have been previously observed in early childhood, early adolescence and adults (Bonmassar et al., 2020; Marois & Vachon, 2018; Ríos-López et al., 2022; Wetzel et al., 2016; Widmann et al., 2018). Thus, our results contribute to the growing body of literature linking involuntary attention to changes in pupil size (Eckstein et al., 2017; Marois et al., 2020). Similar deviant-related amplitudes in PDR in all age groups could be interpreted as a similar transient increase in arousal during deviant trials in all age groups.

Increased PDR latency indicates longer cognitive processing (Zekveld et al., 2011). The delayed PDR in response to deviant sounds observed across all age groups suggests a prolonged processing of deviant sounds probably due to attentional orienting in all groups. The lack of age differences in the deviant-related PDR, while simultaneously significant age effects on behavioral distraction were observed, indicates that behavioral and pupil measures reflect partly distinct aspects of attentional distraction.

###### 2.3.2 Age effects on executive attention

PDR amplitude and latency were also increased in response to Go as opposed to Nogo sounds in all groups. When comparing Go and Nogo trials, it is important to point out, that the motor response has a considerable impact on pupil dilation (e.g. Hupé et al., 2009). Larger PDR amplitude to Go compared to Nogo trials can be attributed to the additional engagement of motor processes, but also of executive attention processes, in response to the Go sounds. The absence of an age-difference in the Go-Nogo effect on PDR amplitude could indicate a similar increase in sound-related arousal in the younger age groups compared to adults. The decreasing PDR latency with age could at least partly be related to delayed button responses (i.e. larger reaction times) in teens and children and point to differences in target processing speed between groups (Zekveld et al., 2011). The Go/Nogo difference of PDR latency was smaller in adults compared to both teens and children; while it was similar in teens and children. This indicates comparable immature executive attention capacities in children and teens. This protracted maturation of executive attention aligns with the maturation of the frontal lobes, which undergo considerable structural and functional changes well beyond early childhood but mainly during adolescence and puberty (Durston & Casey, 2006; Gogtay et al., 2004; Sowell et al., 2003; Tamm et al., 2002; Toga et al., 2006).

Our results demonstrate that the pupil is a sensitive tool to measure involuntary attention as well as executive attention abilities in young children.

#### 3 ERPs (RIDE decomposition)

**3.1 Methods**

#### Residue iteration decomposition (RIDE). The changes in brain electrical potential evoked by an event consist of a series of deflections (components) whose latencies at the single trial level are either time-locked to stimulus onset, or time-locked to response onset, or have variable latencies and are assumed to reflect aspects of sensory and cognitive processing. Averaging of single trial ERPs locked to stimulus onset only, as typically performed, will pronounce the stimulus-locked components but attenuate and smear the response-locked and variable latency contributions. However, many relevant cognitive processes may occur at variable latency or response-locked (for a more detailed introduction and discussion see Ouyang et al., 2013, 2015). To dissociate the individual average ERPs related to sensory processing (S-component; locked to stimulus onset), to response execution and motor processing (R-component; locked to button press), and a central component with variable latency (C-component; typically in between sensory and motor processing), the residue iteration decomposition method has been developed (RIDE Matlab toolbox; Ouyang et al., 2015) and applied here. Many higher-level cognitive processes, such as conflict resolution and response inhibition are thought to be reflected in the C-component (Ouyang et al., 2015), so we focused our analysis on this component. The time windows used for the RIDE decomposition were 0 to 500 ms for the S-component, 100 to 900 ms for the C-component (both relative to stimulus onset), and −300 to +300 ms for the R-component (relative to response latency; all values reflect the defaults suggested by the toolbox manual and Ouyang et al., 2015). The Nogo-Go difference of C-component was calculated for each sound type and group.

**3.2 Results**

###### 3.2.1 Variability of the RIDE C-component

The variability of the RIDE C-component latency decreased with age (Ch: 112 ms; Te: 108 ms; Ad: 95 ms). The Bayesian ANOVA preferred the model including the sound type, target and age group main effects and the sound type by target and target by age group interaction effects (BF_10_=7.68×10^5^). The data provided moderate evidence against the sound type by age group (BF_10_=0.222) and sound type by age group by target interaction effects (BF_10_=0.297) The C-component variability was higher for Go trials in standards and higher for Nogo trials in deviants. The data only provide anecdotal evidence for both effects (Sta Go vs Nogo: BF_10_=0.782; Dev: BF_10_=2.38). Similarly, the C-component variability was higher for Nogo than Go trials in children, similar in teens and higher for Go than Nogo trials in adults. Again, the data only provide anecdotal evidence for any of these effects (Ch Go vs Nogo: BF_10_=1.69; Te: BF_10_=0.199; Te: BF_10_=1.33). The data provided anecdotal evidence against a difference in the C-component variability between children and teens (BF_10_=0.557) but strong evidence for a difference between children and adults (BF_10_=2.08×10^4^) and teens and adults (BF_10_=728). Therefore, we consider the application of the RIDE decomposition as necessary to compare the ERP data between age groups.

###### 3.2.2 P3a (S-component)

We observed more positive S-component amplitudes around 250-350 ms over central areas in response to deviant stimuli compared to standard stimuli. We consider this difference potential which is best visible in the deviant minus standard difference wave (Figure S3) as the P3a. The P3a peak latency was 344 ms for children, 312 ms for teens, and 264 ms for adults with peak amplitudes at Cz electrode location in all age groups, on the difference deviant minus standard difference (Figure S3).

**Latency.** The Bayesian ANOVA on the P3a peak latency preferred the null model. The data provided moderate evidence against a P3a latency difference between Go and Nogo stimuli (BF_incl_=0.166), moderate evidence against a P3a latency difference between age groups (BF_incl_=0.130), and strong evidence against a sound type by target by age group interaction effect (BF_incl_=0.098).

**Amplitude.** We computed the mean amplitudes in (80 ms duration) time windows centered on the peak of the P3a per age group. The Bayesian ANOVA on the P3a amplitudes preferred the model including the sound type and age group main effects (BF_10_=8.46×10^13^). The data provided strong evidence for a sound type main effect (BF_incl_=7.69×10^12^) and for a group main effect (BF_incl_= 11.2), and anecdotal evidence against both a sound type by age group (BF_incl_=0.865) and a sound type by target by age group interaction effects (BF_incl_=0.384). As the evidence against the sound type by age group interaction was rather inconclusive we wanted to verify P3a elicitation separately for age groups. The data provided moderate to strong evidence for an effect of sound type in all age groups (Sta < Dev in Ch: BF_10_=5.3, Te: BF_10_=5.50×10^5^, Ad: BF_10_=6.15×10^5^).

In sum, we observed a P3a for both Go and Nogo stimuli in all age groups, indicating comparable attentional orienting and arousal in all age groups during the present Go/Nogo task.

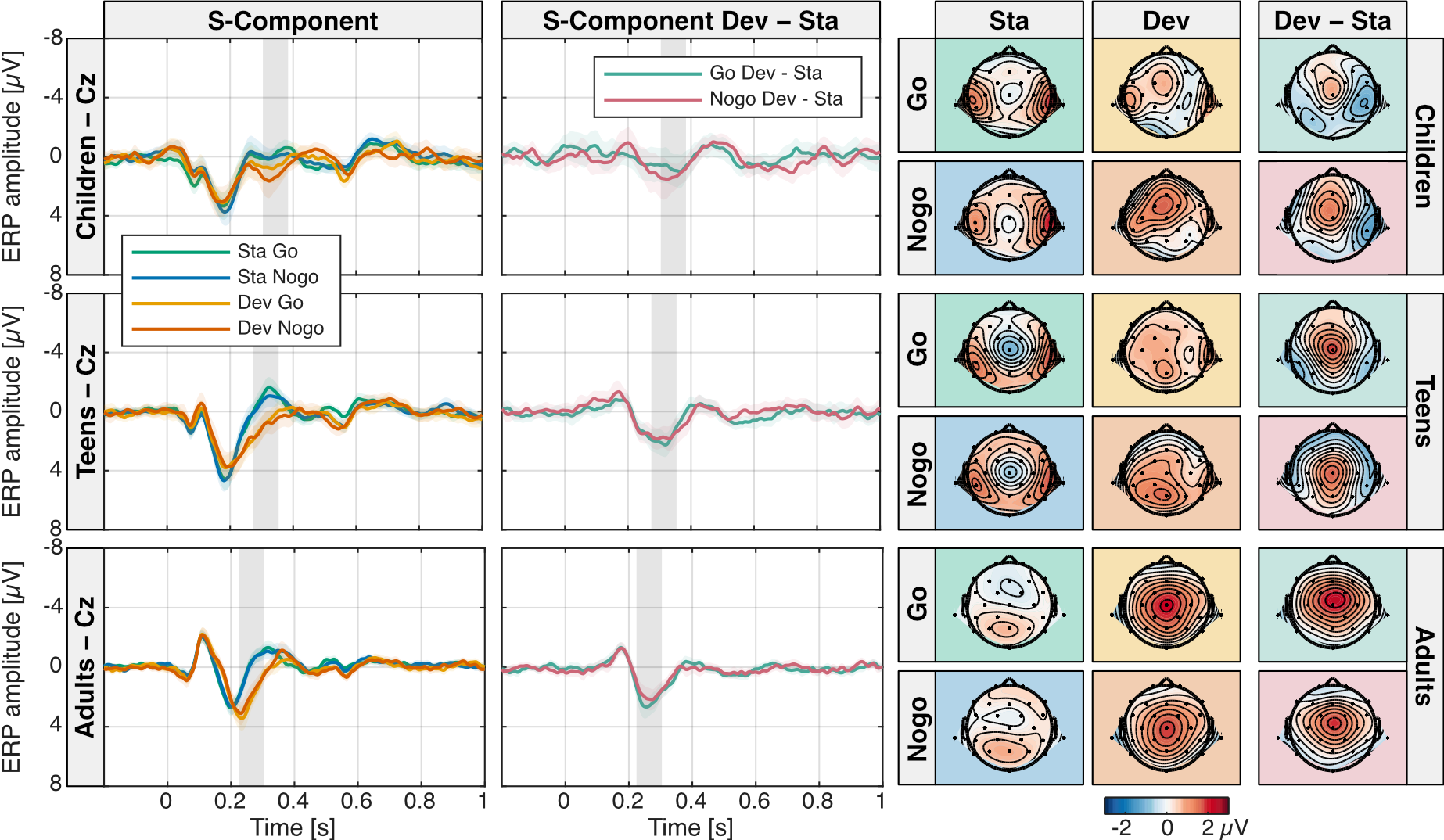

**Figure S3: P3a in the RIDE S-component.** A central P3a was observed in the deviant minus standard difference wave for Go and Nogo stimuli in all age groups. Left: RIDE S-component standard and deviant, Go and Nogo ERP waves at electrode location Cz in children, teens and adults. Shaded areas indicate 95% confidence intervals. Middle: Corresponding deviant minus standard difference waves of the S-component for both target conditions (standard: light blue, deviant: pink) and age groups. The grey bar indicates the P3a analysis time window. Right: RIDE S-component Go and Nogo, standard and deviant ERP topographies in the P3a time window and deviant minus standard difference topographies for all age groups.

###### 3.2.3 P3b (C-Component)

We observed a P3b component between 550 and 650 ms over parietal areas in response to any stimulus type. P3b amplitude was higher for Go compared to Nogo, higher for deviants compared to standards, and also higher in children and teens compared to adults (Figure S4). The P3b peak latency decreased with age between children and teens and was longer for deviants compared to standards (sta vs dev in Ch: 616 vs 652 ms, Te: 560 vs 608 ms, Ad: 552 vs 588 ms).

**Latency.** The Bayesian ANOVA on the P3b peak latency preferred the model including the sound type and age group main effects only (BF_10_=1563). The data provided moderate to strong evidence against a main effect of target, all possible two-way and the three-way interaction (target: BF_incl_=0.131; sound type × target: BF_incl_=0.169; sound type × age group: BF_incl_=0.080; target × age group: BF_incl_=0.101; sound type × target × age group: BF_incl_=0.168). The data provided strong evidence for a difference in P3b latency between children and teens (BF_10_=318) and children and adults (BF_10_=37.7) and anecdotal evidence against a difference between teens and adults (BF_10_=0.397).

**Amplitude.** We computed the mean amplitudes in (80 ms duration) time windows centered on the peak of the P3b per age group and sound type. The Bayesian ANOVA on the P3b amplitudes preferred the model including the sound type, target and age group main effects and the sound type by age group interaction effect (BF_10_=5.74×10^11^). The data provided strong evidence for age (BF_incl_=294), sound type (BF_incl_=3497) and target main effects (BF_incl_=2.93×10^5^), moderate evidence against sound type by target (BF_incl_=0.206) and target by age group interaction effects (BF_incl_=0.109) and anecdotal evidence against a sound type by target by age group interaction effect (BF_incl_=0.540).

The data provided anecdotal and strong evidence for higher P3b amplitudes for deviants compared to standards in children (BF_incl_=2.18) and teens (BF_incl_=4979), respectively and moderate evidence against such a difference in adults (BF_incl_=0.280). The data provided moderate evidence that P3b amplitudes were similar rather than different between children and teens (BF_incl_=0.260), and strong evidence that P3b amplitudes were smaller in adults (Ch vs Ad: BF_incl_=42.7; Te vs Ad: BF_incl_=2116).

In sum, P3b latency decreased with age and was longer for deviants than standards; P3b amplitudes were larger for Go than Nogo, larger for deviants than standards (except in adults) and smaller in adults.

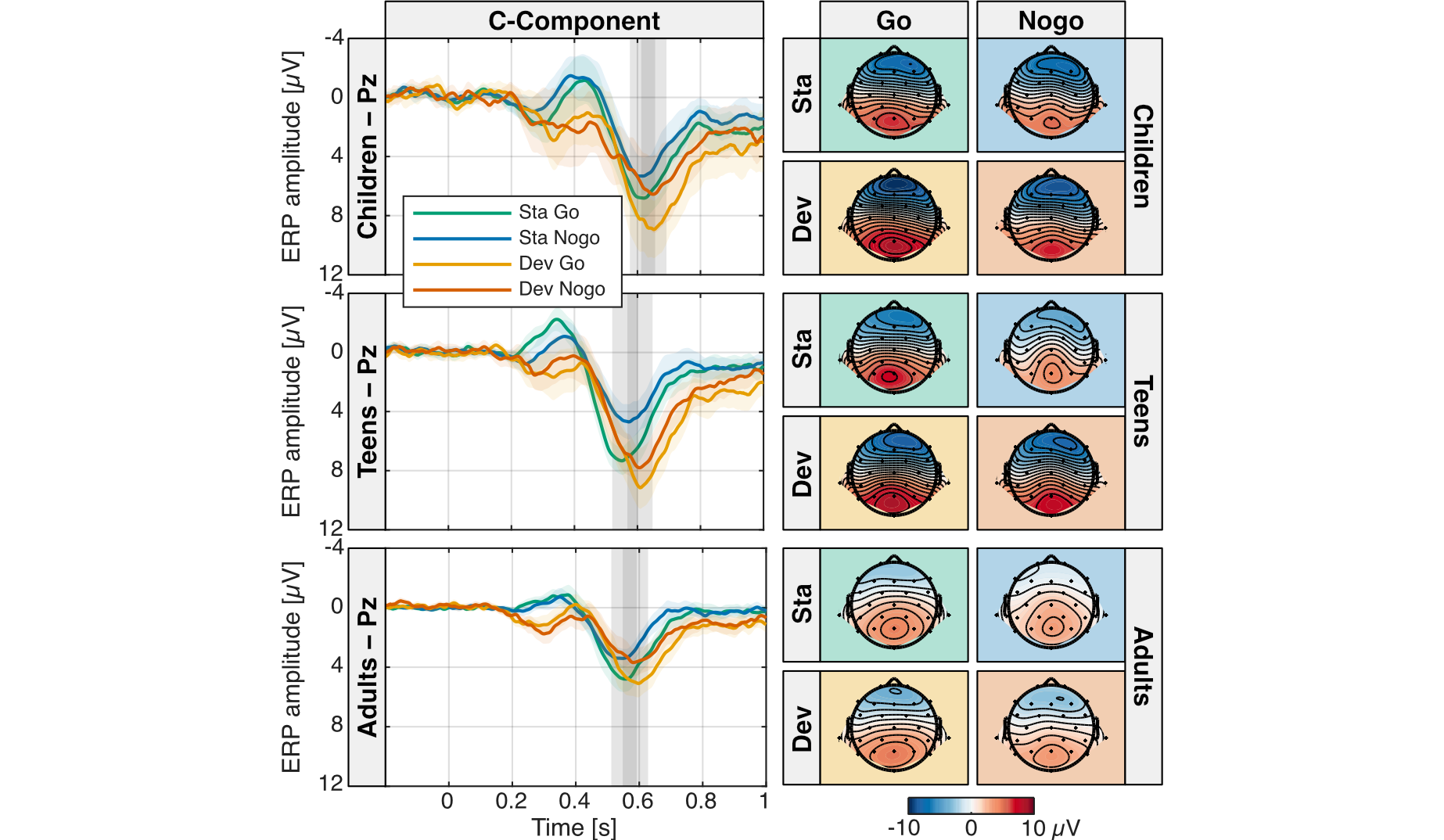

**Figure S4: P3b in the RIDE C-component.** A parietal P3b was observed in all conditions and age groups. P3b latency was longer for deviants than standards. P3b amplitude was higher for Go than Nogo and higher for deviants than standards (except in adults) and smaller in adults. Left: RIDE C-component standard and deviant, Go and Nogo ERP waves at electrode location Pz in children, teens and adults. Shaded areas indicate 95% confidence intervals. The grey bar indicates the P3b analysis time window. Right: RIDE C-component Go and Nogo, standard and deviant ERP topographies in the P3b time window.

##### **3.3 Discussion**

###### 3.3.1 Age effects on the P3b

As expected, Go compared to Nogo trials evoked larger amplitudes of the parietal P3b in all age groups. Remarkable age differences in the processing of standard and deviant trials were observed. Deviant trials evoked significant larger amplitudes compared to standard trials in teens, and with less statistical evidence in children. However, there was statistical evidence against a difference in amplitude between standard and deviant trials in adults, in contradiction with many studies with adults showing larger amplitude of the P3b as the temporal probability of a target stimulus decreases (Polich, 2007). Here we observed this pattern in children and especially in teens, who exhibit a larger amplitude in infrequent deviant trials than in frequent standard trials. Various hypotheses have been proposed regarding the P3b underlying processes (for a review, see Polich, 2007). In addition to uncertainty, one influencing factor is the allocation of attentional resources (Kahneman, 1973; Polich, 2007). The present results suggest that adolescents (and children) allocate more resources to task-irrelevant deviant than standard sounds at this stage of cognitive processing, whereas adults similarly processed deviant and standard sounds. The latter is more resource-efficient because sound features defining deviant sounds are not relevant for the task.

Statistics favored the model without an interaction between executive attention and deviant processing (BF_incl_=0.206), that is the distinct processing of standard and deviant sounds in the age groups was statistically similar in Go and Nogo trials. This suggests no more impact of deviant processing on the response inhibition and task performance at this stage of processing.

While mean amplitudes of the P3b in teens were similar to children but different from adults, the latency of the P3b in teens was similar to those of adults but different from children. P3 latency has been assumed to reflect classification speed, namely the time necessary for detection and evaluation of the target stimulus (Polich, 2007). The different developmental pathway for amplitudes and latency of the P3b separates and specifies the development of the respective target-related attention mechanisms throughout childhood.

#### 4 ERPs

###### **4.1 Results**

###### 4.1.1 P3a (ERPs)

**Latency**. In the ERPs the P3a latency decreased with age from 352 ms in children, 324 ms in teens to 264 ms in adults. In contrast to the corresponding RIDE S-component analysis, the Bayesian ANOVA on the ERP P3a latencies preferred the model including the group main effect (BF_10_=6.144). The data provided moderate evidence against a target main effect (BF_incl_=0.156) and the target by age group interaction effect (BF_incl_=0.122). The data provided anecdotal evidence against a difference in P3a latency between children and teens (BF_10_=0.371), moderate evidence for a higher latency in children than adults (BF_10_=7.38) and strong evidence for a higher latency in teens than adults (BF_10_=6549).

**Amplitude**. We computed the mean amplitudes in (80 ms duration) time windows centered on the peak of the P3a in the deviant minus standard difference waves per age group. The Bayesian ANOVA on the P3a ERP amplitudes replicates the results obtained for the RIDE S-component. It also preferred the model including the sound type and age group main effects (BF_10_=3.11×10^14^). The data provided strong evidence for a sound type main effect (BF_incl_=2.98×10^12^) and anecdotal evidence against both a sound type by age group (BF_incl_=0.989) and a sound type by target by age group interaction effects (BF_incl_=0.314). As the evidence against the sound type by age group interaction was rather inconclusive we wanted to verify P3a elicitation separately for age groups. The data provided moderate to strong evidence for an effect of sound type in all age groups (Sta < Dev in Ch: BF_10_=5.37, Te: BF_10_=1.38×10^5^, Ad: BF_10_=6.15×10^5^). In sum, we replicated the result of a P3a in all age groups.

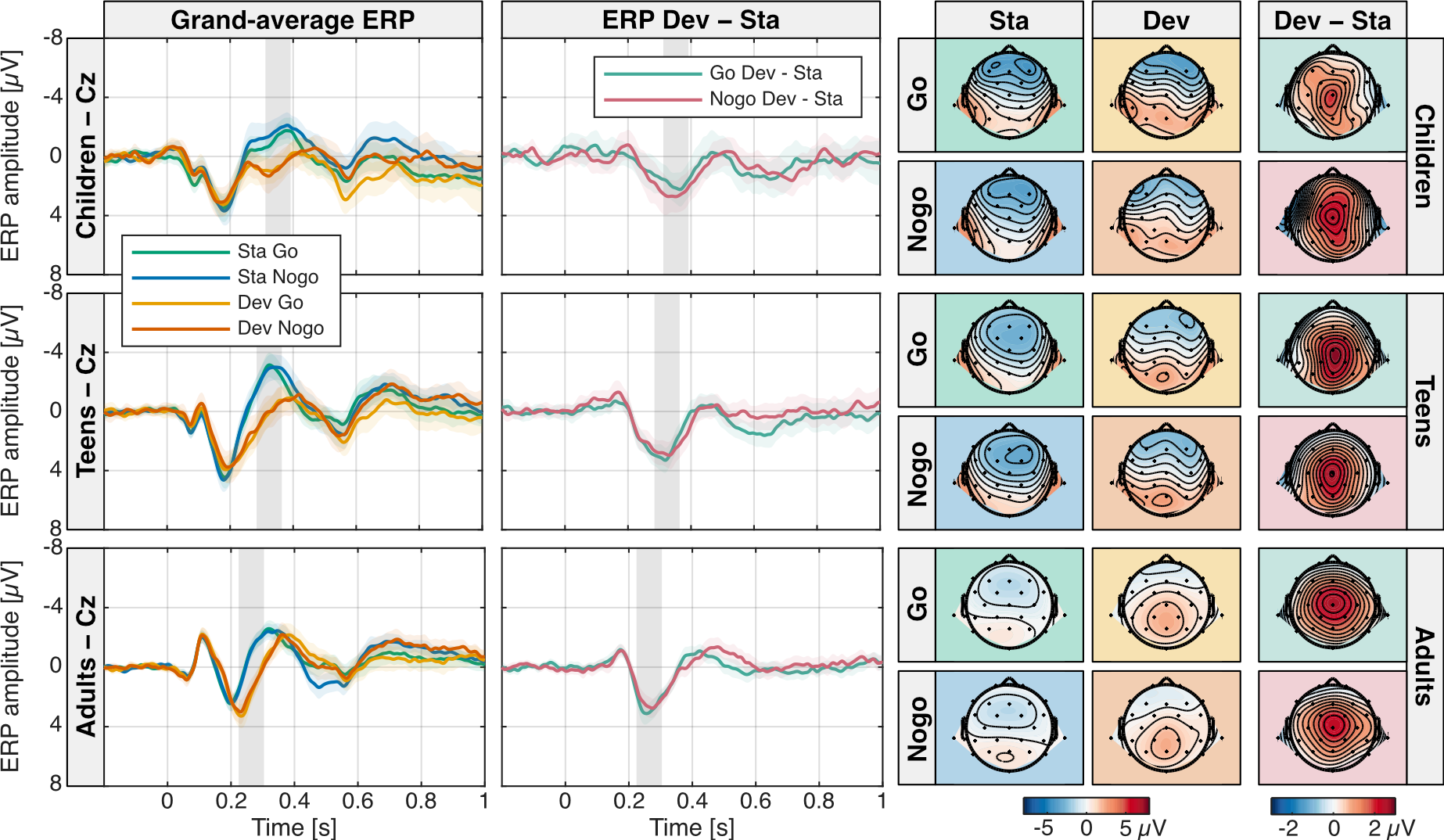

**Figure S5: P3a.** A central P3a was observed in the deviant minus standard difference wave for Go and Nogo stimuli in all age groups. Left: Standard and deviant, Go and Nogo grand-average ERP waves at electrode location Cz in children, teens and adults. Shaded areas indicate 95% confidence intervals. Middle: Corresponding deviant minus standard difference waves of the grand-average ERPs for both target conditions (standard: light blue, deviant: pink) and age groups. The grey bar indicates the P3a analysis time window. Right: Grand-average ERP Go and Nogo, standard and deviant ERP topographies in the P3a time window and deviant minus standard difference topographies for all age groups.

###### 4.1.2 Nogo-N2 effect (ERPs)

**Latency**. The Nogo-N2 effect latency analysis on the ERP data did not replicate the corresponding RIDE C-component analysis. The Bayesian ANOVA on the standard Nogo-N2 effect latency preferred the null model. The data provided anecdotal evidence against a main effect of age group (BF_incl_=0.357). We consider the absence of an age group latency effect in the ERP data to be likely due to the higher noise level of the ERP (see for example the multiple peaks in teens in the time window of the N2). To avoid spurious Nogo-N2 effect amplitude effects we still computed the mean amplitudes in (40 ms duration) time windows centered on the peak of the Nogo-N2 effect in standards per age group (476 ms for children, 412 ms for teens, and 372 ms for adults) despite the absent age group latency effect.

**Amplitude**. The Bayesian ANOVA on the Nogo-N2 effect amplitudes replicates the results obtained for the RIDE C-component (except for an age group main effect). It preferred the model including the sound type and target main effects and the sound type by target interaction effects (BF_10_=100.5). The data provided moderate evidence against a target by age group (BF_incl_=0.096) and a sound type by target by age group interaction effect (BF_incl_=0.247). There was strong evidence for a Nogo-N2 effect in standards (Nogo vs Go: BF_10_=6250) but moderate evidence against a Nogo-N2 effect in deviants (BF_10_=0.115). In sum, we replicated the result of a Nogo-N2 effect for standards but not for deviants in all age groups in the ERP data.

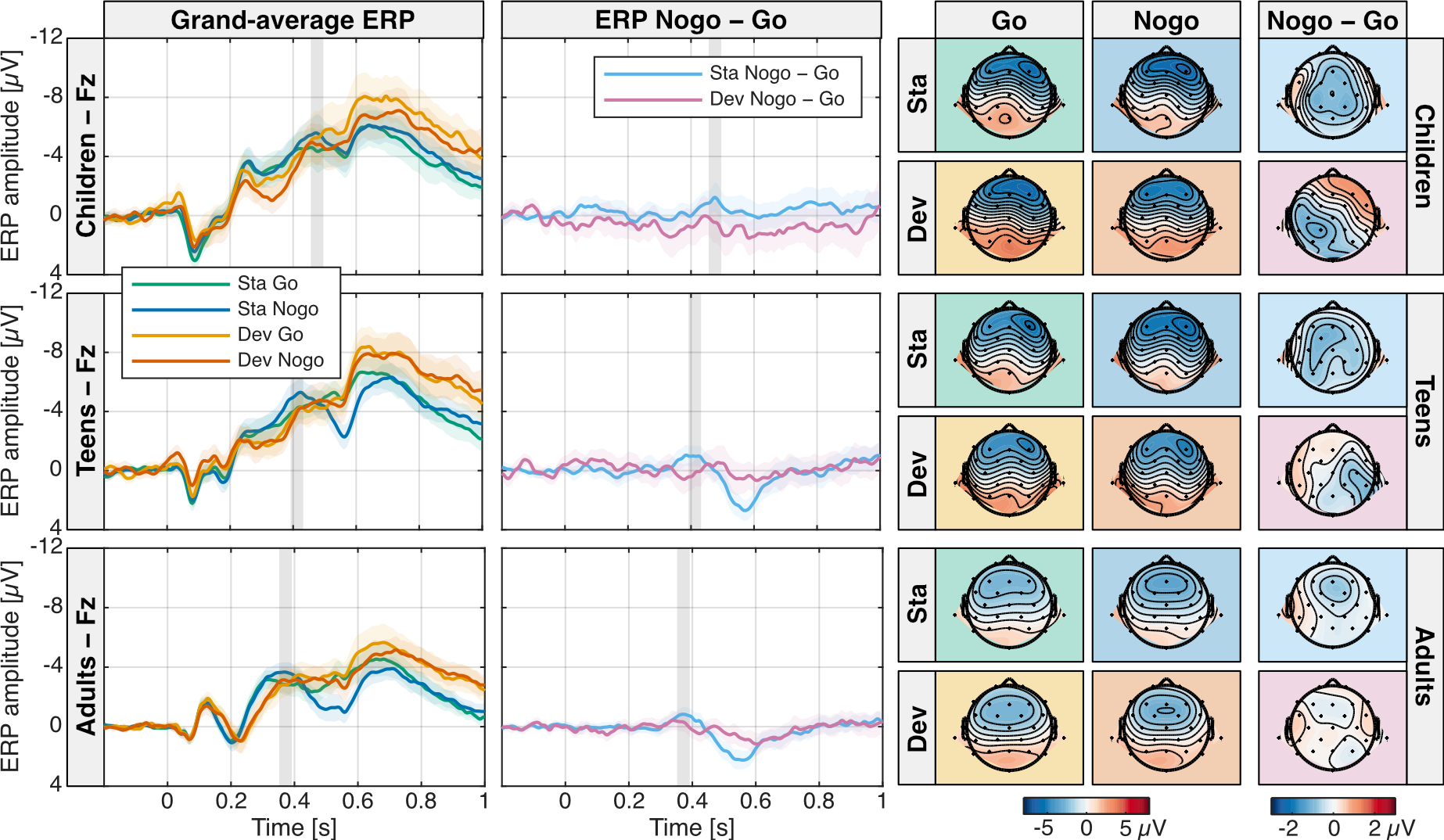

**Figure S6: Nogo-N2 effect.** A frontal Nogo-N2 was observed in the Nogo minus go difference wave for standards but not for deviants in all age groups. Left: Standard and deviant, Go and Nogo grand-average ERP waves at electrode location Fz in children, teens and adults. Shaded areas indicate 95% confidence intervals. Middle: Corresponding Nogo minus Go difference waves of the grand-average ERPs for both sound type conditions (standard: light blue, deviant: pink) and age groups. The grey bar indicates the Nogo-N2 analysis time window. Right: Grand-average ERP standard and deviant, Go and Nogo ERP topographies in the Nogo-N2 time window and Nogo minus Go difference topographies for all age groups.

###### 4.1.3 Nogo-P3 effect (ERPs)

**Latency**. As in the RIDE C-component analysis, the data provided moderate evidence against a latency difference in the Nogo-P3 effect in standards between teens and adults (BF_10_=0.306) where a Nogo-P3 effect peak could be identified (Te: 572 ms; Ad: 560 ms).

**Amplitude**. We computed the mean amplitudes in (80 ms duration) time windows centered on the peak of the Nogo-P3 effect in standards in teens and adults and the most positive peak in the corresponding time range at 616 ms in children. The Bayesian ANOVA on the Nogo-P3 effect amplitudes replicates the results obtained for the RIDE C-component. It preferred the same model including the sound type, target and age group main effects and the sound type by task and task by age group interaction effects (BF_10_=1.03×10^12^). The data provided moderate evidence against a sound type by age group (BF_incl_=0.097) and anecdotal evidence for a sound type by target by age group interaction effect (BF_incl_=1.7). There was strong evidence for a Nogo-P3 effect in standards (Nogo vs Go: BF_10_=1726) but moderate evidence against a Nogo-P3 effect in deviants (BF_10_=0.189). There was moderate evidence against a Nogo-P3 effect in children (Nogo vs Go: BF_10_=0.195) but strong evidence for a Nogo-P3 effect in teens (BF_10_=88.3) and adults (BF_10_=10.4). In sum, we replicated the result of a Nogo-P3 effect for standards in teens and adults but not in children and not for deviants in any age group in the ERP data.

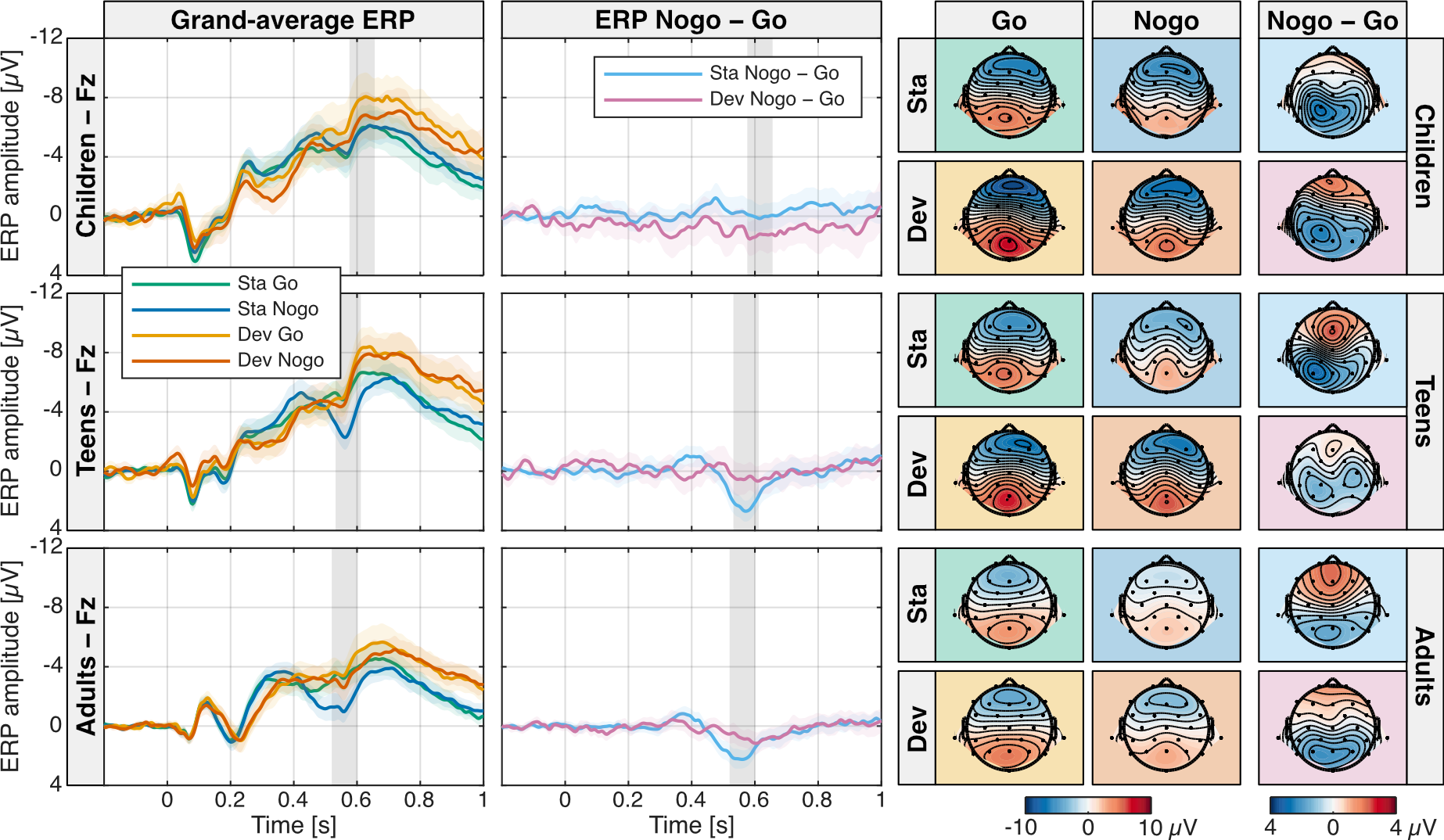

**Figure S7: Nogo-P3 effect.** A frontal Nogo-P3 was observed in the Nogo minus go difference wave for standards but not for deviants in teens and adults but not in children. Left: Standard and deviant, Go and Nogo grand-average ERP waves at electrode location Fz in children, teens and adults. Shaded areas indicate 95% confidence intervals. Middle: Corresponding Nogo minus Go difference waves of the grand-average ERPs for both sound type conditions (standard: light blue, deviant: pink) and age groups. The grey bar indicates the Nogo-P3 analysis time window. Right: Grand-average ERP standard and deviant, Go and Nogo ERP topographies in the Nogo-P3 time window and Nogo minus Go difference topographies for all age groups.

###### 4.1.4 P3b (ERPs)

**Latency**. The P3b latency analysis on the ERP data did not replicate the corresponding RIDE C-component analysis. The Bayesian ANOVA on the standard P3b latency preferred the null model. The data provided anecdotal evidence against a main effect of age group (BF_incl_=0.818) and moderate evidence against all other main and interaction effects (sound type: BF_incl_=0.236; target: BF_incl_=0.130; sound type × target: BF_incl_=0.150; sound type × age group: BF_incl_=0.076; target × age group: BF_incl_=0.087; sound type × target × age group: BF_incl_=0.149). We consider the absence of a sound type or age group latency effect in the ERP data to be likely due to the higher noise level of the ERP (we observed multiple potential P3b peaks in all age groups).

**Amplitude**. To avoid spurious P3b amplitude effects we still computed the mean amplitudes in (120 ms duration) time windows centered on the peak of the P3b averaged over conditions per age group (684 ms for children, 584 ms for teens, and 576 ms for adults) despite the absent age group latency effect. The Bayesian ANOVA on the P3b ERP amplitudes replicates the results obtained for the RIDE C-component. It also preferred the model including the sound type, target and age group main effects and the sound type by age group interaction effect (BF_10_=1.69×10^11^). The data provided strong evidence for sound type (BF_incl_=4.89×10^4^) and target main effects (BF_incl_=2.24×10^4^), moderate evidence against sound type by target (BF_incl_=0.241) and target by age group interaction effects (BF_incl_=0.095) and anecdotal evidence against a sound type by task by age group interaction effect (BF_incl_=0.582). The data provided strong evidence for higher P3b amplitudes for deviants compared to standards in children (BF_incl_=29.3) and teens (BF_incl_=225), respectively and anecdotal evidence against such a difference in adults (BF_incl_=0.563). The data provided anecdotal evidence that P3b amplitudes were similar rather than different between children and teens (BF_incl_=0.368), and moderate to strong evidence that P3b amplitudes were smaller in adults (Ch < Ad: BF_incl_=6.77; Te < Ad: BF_incl_=889). In sum, we replicated the results of higher P3b amplitudes for Go than Nogo, and also deviants than standards (except in adults), and smaller P3b amplitudes in adults in the ERP data.

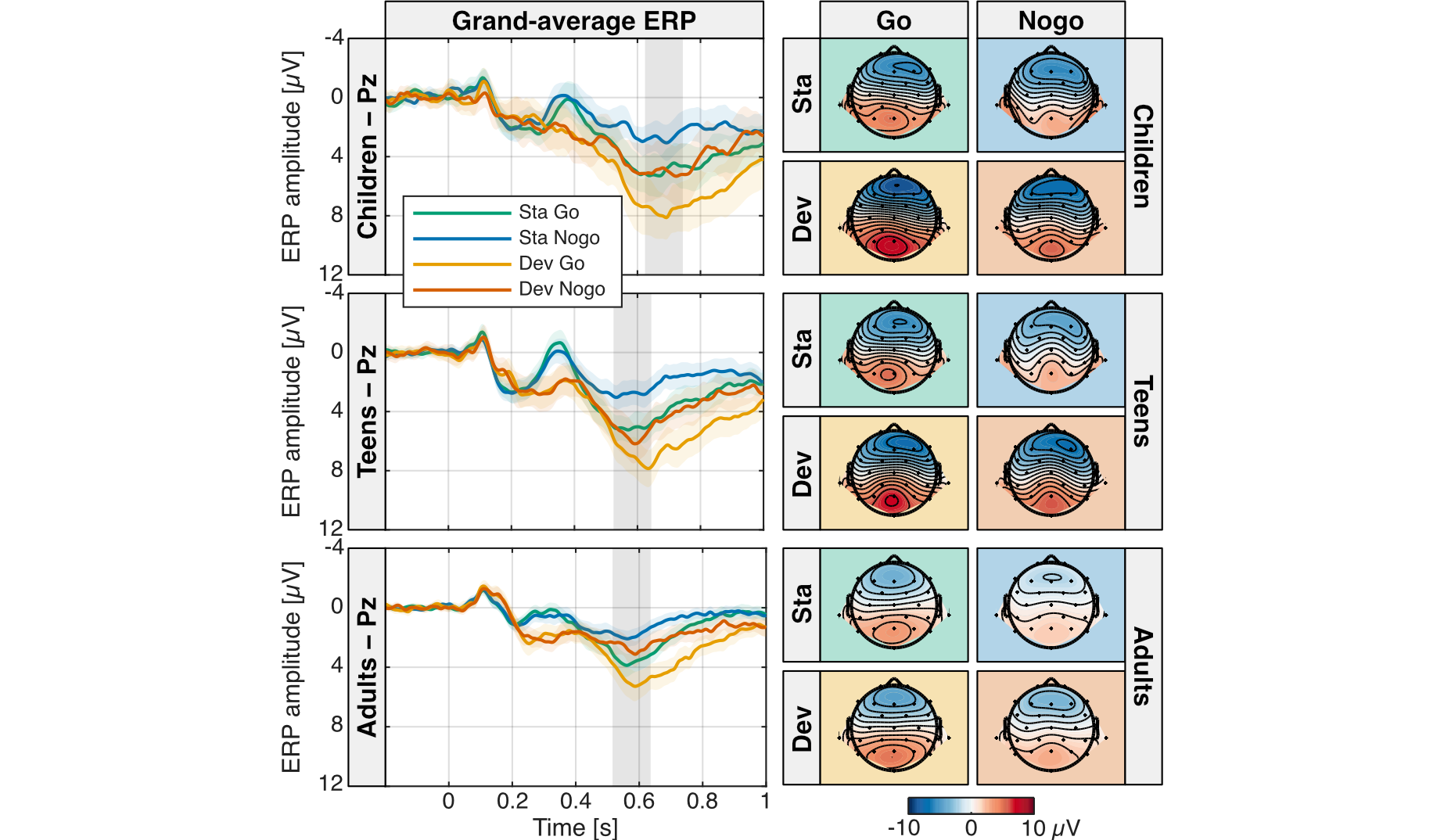

**Figure S8: P3b.** A parietal P3b was observed in all conditions and age groups. P3b amplitude was higher for Go than Nogo and higher for deviants than standards (except in adults) and smaller in adults. Left: Standard and deviant, Go and Nogo grand-average ERP waves at electrode location Pz in children, teens and adults. Shaded areas indicate 95% confidence intervals. The grey bar indicates the P3b analysis time window. Right: Grand-average ERP Go and Nogo, standard and deviant ERP topographies in the P3b time window.
